# Effects of persistent sodium current blockade in respiratory circuits depend on the pharmacological mechanism of action and network dynamics

**DOI:** 10.1101/575688

**Authors:** Ryan S. Phillips, Jonathan E. Rubin

## Abstract

The mechanism(s) of action of most commonly used pharmacological blockers of voltage-gated ion channels are well understood; however, this knowledge is rarely considered when interpreting experimental data. Effects of blockade are often assumed to be equivalent, regardless of the mechanism of the blocker involved. Using computer simulations, we demonstrate that this assumption may not always be correct. We simulate the blockade of a persistent sodium current (*I*_*NaP*_), proposed to underlie rhythm generation in pre-Bötzinger complex (pre-BötC) respiratory neurons, via two distinct pharmacological mechanisms: (1) pore obstruction mediated by tetrodotoxin and (2) altered inactivation dynamics mediated by riluzole. The reported effects of experimental application of tetrodotoxin and riluzole in respiratory circuits are diverse and seemingly contradictory and have led to considerable debate within the field as to the specific role of *I*_*NaP*_ in respiratory circuits. The results of our simulations match a wide array of experimental data spanning from the level of isolated pre-BötC neurons to the level of the intact respiratory network and also generate a series of experimentally testable predictions. Specifically, in this study we: (1) provide a mechanistic explanation for seemingly contradictory experimental results from in vitro studies of *I*_*NaP*_ block, (2) show that the effects of *I*_*NaP*_ block in in vitro preparations are not necessarily equivalent to those in more intact preparations, (3) demonstrate and explain why riluzole application may fail to effectively block *I*_*NaP*_ in the intact respiratory network, and (4) derive the prediction that effective block of *I*_*NaP*_ by low concentration tetrodotoxin will stop respiratory rhythm generation in the intact respiratory network. These simulations support a critical role for *I*_*NaP*_ in respiratory rhythmogenesis in vivo and illustrate the importance of considering mechanism when interpreting and simulating data relating to pharmacological blockade.

**Author summary:** The application of pharmacological agents that affect transmembrane ionic currents in neurons is a commonly used experimental technique. A simplistic interpretation of experiments involving these agents suggests that antagonist application removes the impacted current and that subsequently observed changes in activity are attributable to the loss of that current’s effects. The more complex reality, however, is that different drugs may have distinct mechanisms of action, some corresponding not to a removal of a current but rather to a changing of its properties. We use computational modeling to explore the implications of the distinct mechanisms associated with two drugs, riluzole and tetrodotoxin, that are often characterized as sodium channel blockers. Through this approach, we offer potential explanations for disparate findings observed in experiments on neural respiratory circuits and show that the experimental results are consistent with a key role for the persistent sodium current in respiratory rhythm generation.

## Introduction

Pharmacological compounds that selectively block voltage-gated ion channels are a fundamental tool in neuroscience. Much of our current theoretical understanding of the roles of various ion channels derives from the interpretation of data from experiments dependent on pharmacological manipulations. The mechanisms of action for many of the most commonly used pharmaceutical blockers of voltage-gated ion channels are relatively well understood and fall into one of three mechanistic categories: (1) pore obstruction, (2) shift in activation/inactivation curves, or, less commonly, (3) alteration of ion selectivity [1, 2]. Despite this knowledge, the specific mechanisms of blockade are rarely considered when interpreting or simulating experimental data. Generally it is assumed that selective blockade of an ion channel has the same functional implication regardless of the mechanism involved. In this theoretical study, we demonstrate ways that this assumption can break down, with different blockade mechanisms differentially impacting neuronal and circuit activity.

To illustrate this idea we simulated blockade of a persistent sodium current (*I*_*NaP*_) in the respiratory pre-Bötzinger complex (pre-BötC) via two commonly used sodium channel blockers with distinct mechanisms of action: tetrodotoxin (TTX) and riluzole (RZ). TTX directly obstructs the Na+ pore [3], whereas RZ shifts *I*_*NaP*_ inactivation in the hyperpolarizing direction [4, 5, 6]. At low concentrations, both TTX (*≤* 20 *nM*) and RZ (*≤* 20 *µM*) have been shown to selectively block *I*_*NaP*_ over the fast action-potential generating sodium current (*I*_*Na*_) [7, 8, 9, 10]. A caveat to RZ blockade, however, is that RZ has been shown to inhibit excitatory synaptic transmission at concentrations that affect *I*_*NaP*_ [10, 11].

In respiratory circuits, in vitro blocking studies using TTX or RZ have suggested that *I*_*NaP*_ is critical for intrinsic bursting in pacemaker neurons and network rhythm generation in the isolated pre-BötC [9, 12, 13]. These results have led to the hypothesis that *I*_*NaP*_ may be a necessary component of rhythmogenesis in respiratory circuits [14, 15, 16]. This hypothesis, however, has fallen out of favor due to the observation that *I*_*NaP*_ block by RZ fails to stop respiratory rhythms in intact preparations [13]. This conclusion is dependent on the assumption that RZ effectively blocks *I*_*NaP*_ in an in vivo setting.

This study predicts that after blockade via RZ, *I*_*NaP*_ can be reactivated by transient hyperpolarizing perturbations due to the specific mechanism of action of RZ. Simulated TTX blockade of *I*_*NaP*_ does not yield this same effect. In intact preparations, the pre-BötC receives strong inhibition during the interburst interval [13, 17, 18, 19, 20], which may allow *I*_*NaP*_ to recover from inactivation even after RZ application.

Consistent with this idea, our simulations of the intact respiratory network predict that RZ will fail to effectively block *I*_*NaP*_, while complete block of *I*_*NaP*_ by experimental application of low concentration TTX (*≤*20 *nM*) within the pre-BötC will abolish the respiratory rhythm in the intact respiratory network. Therefore, the failure of RZ to stop respiratory rhythms in intact experimental preparations is not sufficient to rule out a central role for *I*_*NaP*_ in respiratory rhythm generation, which our study supports.

More generally, these simulations illustrate the importance of considering the mechanism of action when interpreting and simulating experimental data from pharmacological blocking studies.

## Results

To illustrate the difference in effects that can arise through blockade of the same current with pharmacological agents acting through different biophysical mechanisms, we focused on the blockade of *I*_*NaP*_ via RZ and TTX in respiratory neurons of the pre-BötC. Experiments have clearly established the presence of *I*_*NaP*_ in neurons of this type and have suggested that neurons exhibiting *I*_*NaP*_ -dependent intrinsic bursting capabilities play a critical role in rhythm generation in the pre-BötC [9, 12, 14]. Therefore, we first reconstructed a neuron model capable of generating *I*_*NaP*_ -dependent intrinsic bursting [14] For this set of simulations, the model represents a synaptically isolated pre-BötC neuron. Depending on their level of excitability (controlled by a tonic excitatory synaptic conductance parameter *g*_*Tonic*_), these neurons can exhibit three distinct patterns of activity: silent, bursting and tonic spiking (Fig 1 A). In an intrinsic bursting regime in this model, *I*_*NaP*_ is required to depolarize neurons above the spiking threshold. During the spiking within a burst, however, *I*_*NaP*_ begins to inactivate, which leads to hyperpolarization and burst termination. In the subsequent silent period or interburst interval, the membrane potential *V*_*m*_ is hyperpolarized, which allows for the voltage-dependent recovery of *I*_*NaP*_ from inactivation. After sufficient recovery, sub-threshold activation of *I*_*NaP*_ again depolarizes *V*_*m*_ above the spiking threshold, causing burst initiation and completion of one burst cycle (Fig 1 B).

**Fig 1.**
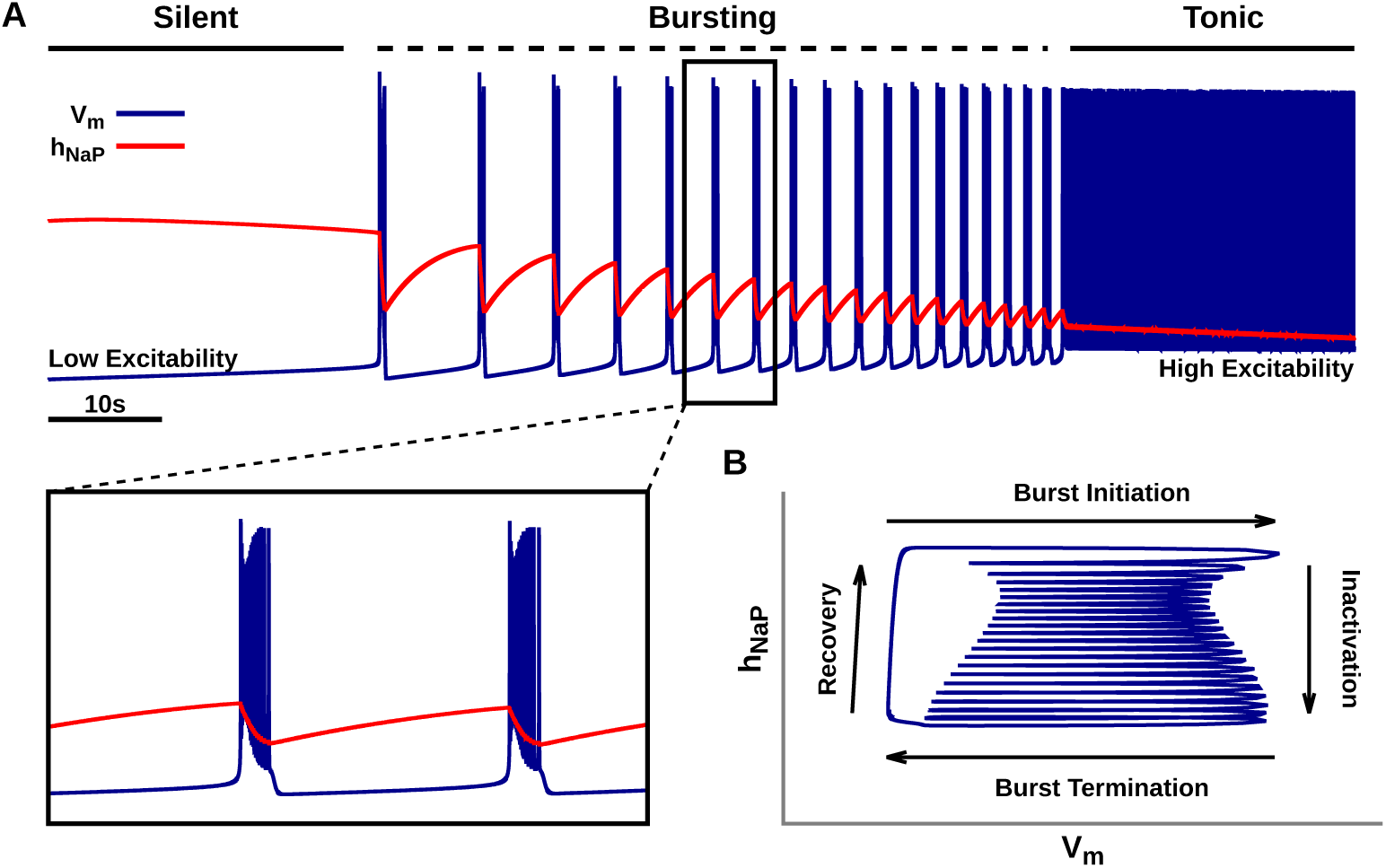
*I*_*NaP*_-dependent intrinsic bursting in a pre-BötC neuron model. (A) Voltage trace illustrating the dependence of simulated neuronal activity patterns on excitability. Inset (bottom left) shows shows burst shape. Excitability was varied by linearly increasing the tonic excitatory synaptic conductance (*g*_*Tonic*_) from 0.25 nS to 4.5 nS over the course of the simulation. (B) Phase plane plot illustrating the relationship between the slow, voltage-dependent *I*_*NaP*_ inactivation variable *h*_*NaP*_ and the membrane potential *V*_*m*_ during the bursting activity displayed in (A). Other model variables are not shown in this view.

### Simulated TTX and RZ blockade of I_NaP_ in steady-state conditions

Bursting in this model is dependent on *I*_*NaP*_, therefore, we first characterized the effects of simulated TTX and RZ blockade on *I*_*NaP*_ under steady-state conditions (Fig 2). Since TTX directly obstructs the pores of sodium channels, TTX blockade was simulated by reducing the conductance *g*_*NaP*_ of *I*_*NaP*_. Since TTX only affects conductance, the steady-state activation and inactivation parameters of *I*_*NaP*_ were not varied (Fig 2 A1). We observed that the steady-state current is reduced proportionally to the decrease of *g*_*NaP*_ (Fig 2 A2). In contrast, RZ shifts voltage-dependent inactivation in the hyperpolarizing direction. Therefore, RZ blockade was simulated by decreasing the half-inactivation parameter *h*_1*/*2_ (Fig 2 B1). We found that, similar to TTX, simulated RZ application effectively blocks *I*_*NaP*_ in steady-state conditions (Fig 2 B2). Interestingly, although TTX and RZ blockade of *I*_*NaP*_ work through distinct mechanisms, the effects on the peak steady-state current are remarkably similar (Fig 2 C).

**Fig 2.**
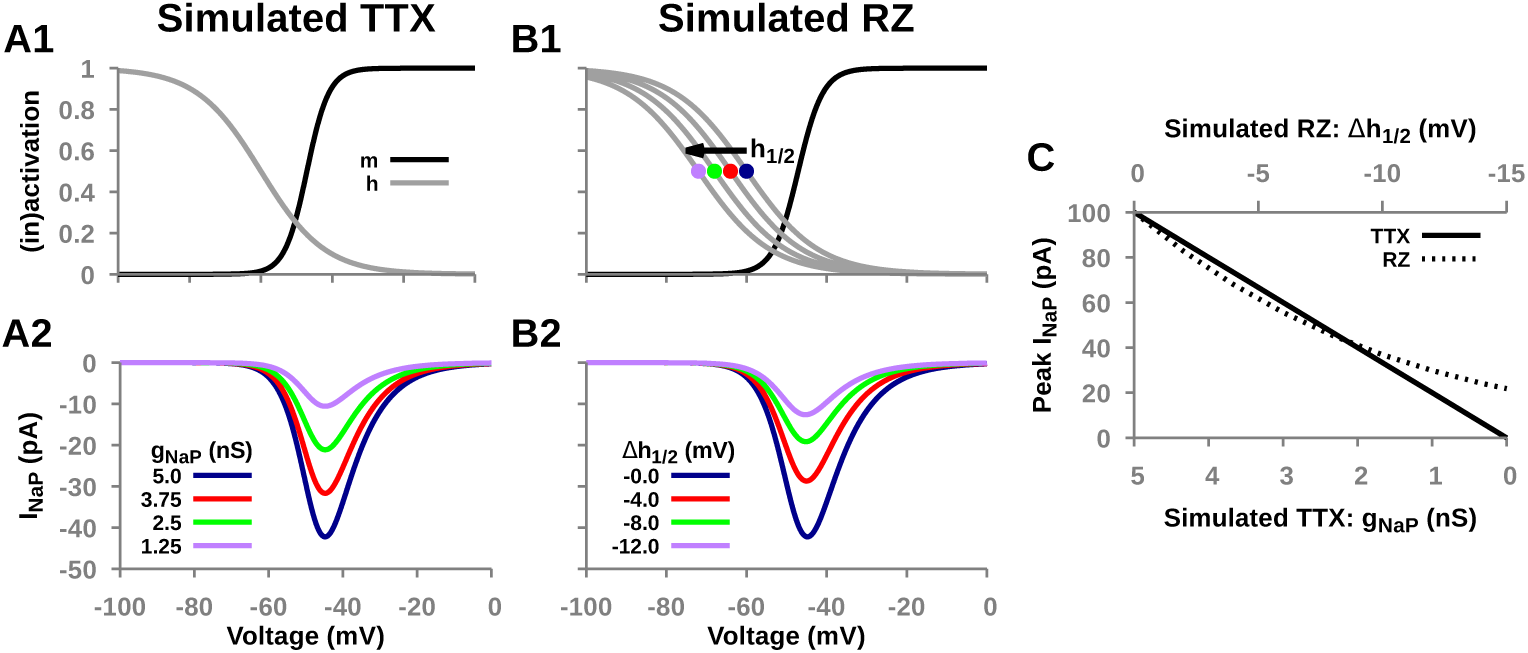
Effects of simulated TTX and RZ blockade on *I*_*NaP*_ under steady-state conditions. (A1 & B1) Effects of simulated TTX and RZ blockade on voltage-dependent steady-state activation (*m*_*∞*_(*V*)) and inactivation (*h*_*∞*_(*V*)) for *I*_*NaP*_. Notice that simulated RZ blockade shifts inactivation in the hyperpolarizing direction and simulated TTX blockade induces no change. (A2 & B2) Effect of simulated TTX and RZ blockade on the I-V curves for *I*_*NaP*_. Notice that simulated TTX and RZ blockade have nearly indistinguishable effects. (C) Peak *I*_*NaP*_ current as a function of the extent of simulated TTX and RZ blockade. TTX and RZ blockade are simulated by reducing *I*_*NaP*_ conductance (*g*_*NaP*_) and shifting its half-inactivation (*h*_1*/*2_) in the hyperpolarizing direction, respectively.

### Simulated TTX and RZ blockade of I_NaP_ abolish intrinsic bursting in an isolated model pre-BötC neuron

The activity pattern exhibited by a neuron capable of *I*_*NaP*_ -dependent bursting is determined by its excitability. Driving the neuron from low to high excitability will drive the transition from silent to bursting and from bursting to tonic spiking regimes of activity. Therefore, we investigated the effects of simulated TTX and RZ blockade of *I*_*NaP*_, represented by reduction of *g*_*NaP*_ or *h*_1*/*2_, respectively, on the intrinsic dynamics of a model pre-BötC neuron over a range of values of tonic synaptic drive conductance (*g*_*Tonic*_), which tunes excitability. We found that simulated TTX and RZ blockade of *I*_*NaP*_ both effectively block intrinsic bursting, either converting bursting dynamics to silence or to tonic spiking, depending in part on the value of *g*_*Tonic*_. Moreover, the effects of these two mechanisms on the shape of the bursting regime in the appropriate 2D parameter space are nearly indistinguishable (Fig 3).

**Fig 3.**
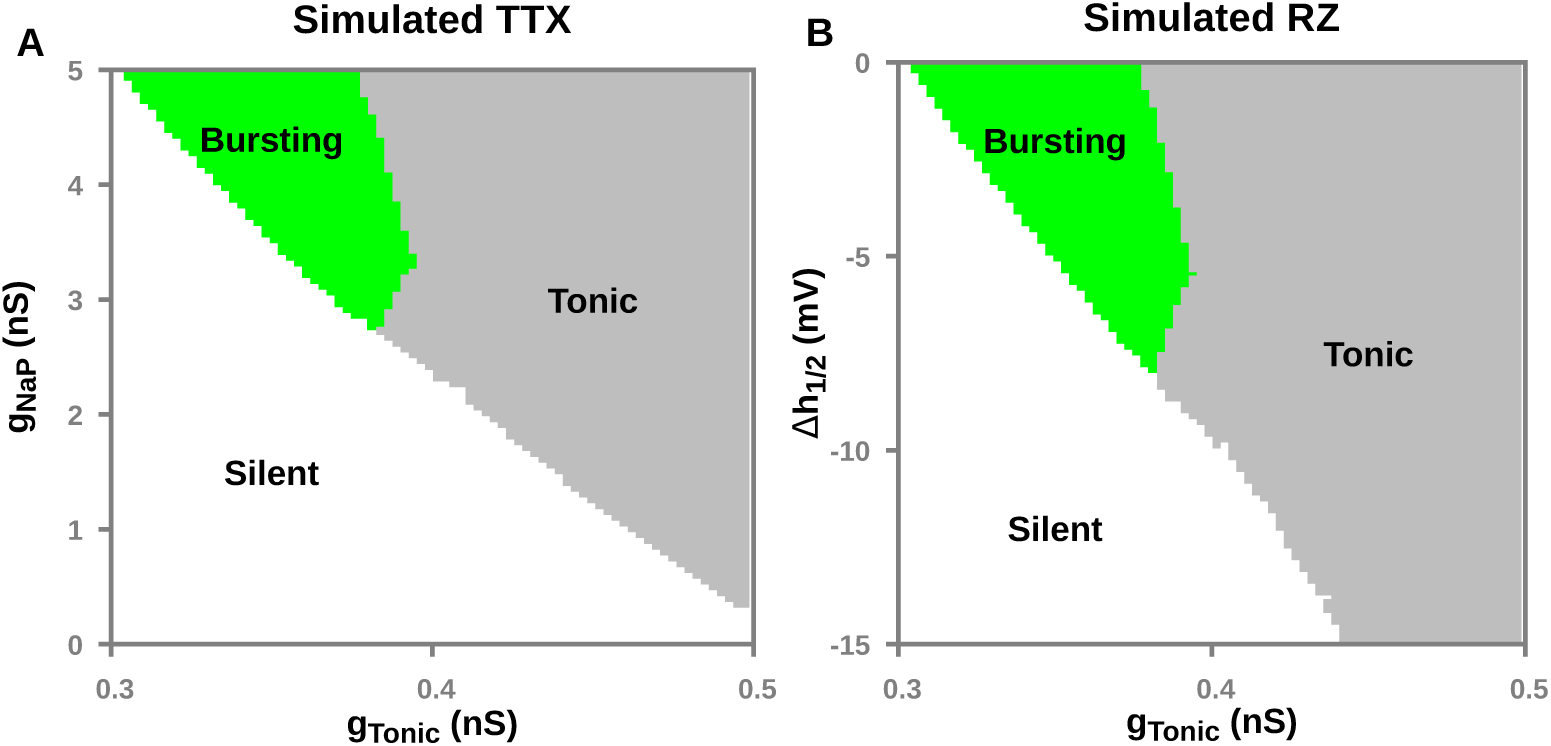
Activity patterns in a model pre-BötC neuron as a function of tonic excitatory synaptic drive (*g*_*Tonic*_) and (A) level of *g*_*NaP*_ and (B) Δ*h*_1*/*2_, which are reduced to simulate TTX and RZ block of *I*_*NaP*_, respectively. Both TTX and RZ effectively abolish intrinsic bursting (green region): bursting does not occur for *g*_*NaP*_ *<* 2.7*nS* nor for Δ*h*_1*/*2_ *< -*8.0*mV*.

Since simulated TTX and RZ blockade have remarkably similar effects on *I*_*NaP*_ and both effectively abolish intrinsic bursting, it is tempting to conclude that although their mechanisms of action are distinct, TTX and RZ are functionally equivalent *I*_*NaP*_ blockers. As we will show next, however, this conclusion is not correct.

### Fast-slow decomposition analysis

Next, to understand how TTX and RZ abolish intrinsic busting, we used fast-slow decomposition analysis (see [21] for review). This method separates the full system into fast and slow subsystems, with the latter represented by *h*_*NaP*_ for the pre-BötC model. Information about the dynamics of the full system can then be inferred from the geometry of the projections of the *V*_*m*_- and *h*_*NaP*_ -nullclines into the (*V*_*m*_, *h*_*NaP*_) phase space. The *V*_*m*_- and *h*_*NaP*_ -nullclines are defined from *dV*_*m*_*/dt* = 0 and *dh*_*NaP*_ */dt* = 0, respectively. The solution to the latter equation is easy to write down as

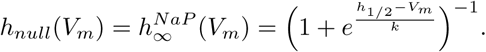

The *V*_*m*_-nullcline is determined numerically to form a manifold consisting of three branches that connect at folds or “knees”, forming an N-shape in the (*V*_*m*_, *h*_*NaP*_)-plane. Intersections between the *V*_*m*_- and *h*_*NaP*_ -nullclines represent equilibrium points of the full system. In the bursting regime, the *h*_*NaP*_ -nullcline intersects the middle branch of the *V*_*m*_-nullcline and forms an unstable equilibrium point that is surrounded by a stable limit cycle.

Simulated TTX and RZ blockade have distinct effects on on the *V*_*m*_- and *h*_*NaP*_ -nullclines. Simulated TTX blockade changes the shape of the *V*_*m*_-nullcline by moving the left and right knees to larger and smaller values of *h*_*NaP*_, respectively, although the right knee is not relevant to the behaviors under study. The former effect corresponds to the increased deinactivation of *I*_*NaP*_ needed for the neuron to activate after *g*_*NaP*_ has been decreased. As *g*_*NaP*_ is lowered, the model remains in the bursting regime until the part of the *V*_*m*_-nullcline near the left knee intersects the *h*_*NaP*_ -nullcline. The bifurcation that results creates a stable equilibrium point that corresponds to the stabilization of a constant, hyperpolarized membrane potential, marking a transition from bursting to silence (Fig 4 B, gray dots). In contrast, simulated RZ blockade only affects the *h*_*NaP*_ -nullcline, inducing a shift in the leftward, hyperpolarizing direction. In this case, as *h*_*NaP*_ is diminished, bursting continues until the *h*_*NaP*_ -nullcline intersects the *V*_*m*_-nullcline near the left knee and forms a stable equilibrium point (Fig 4 C, gray dots).

**Fig 4.**
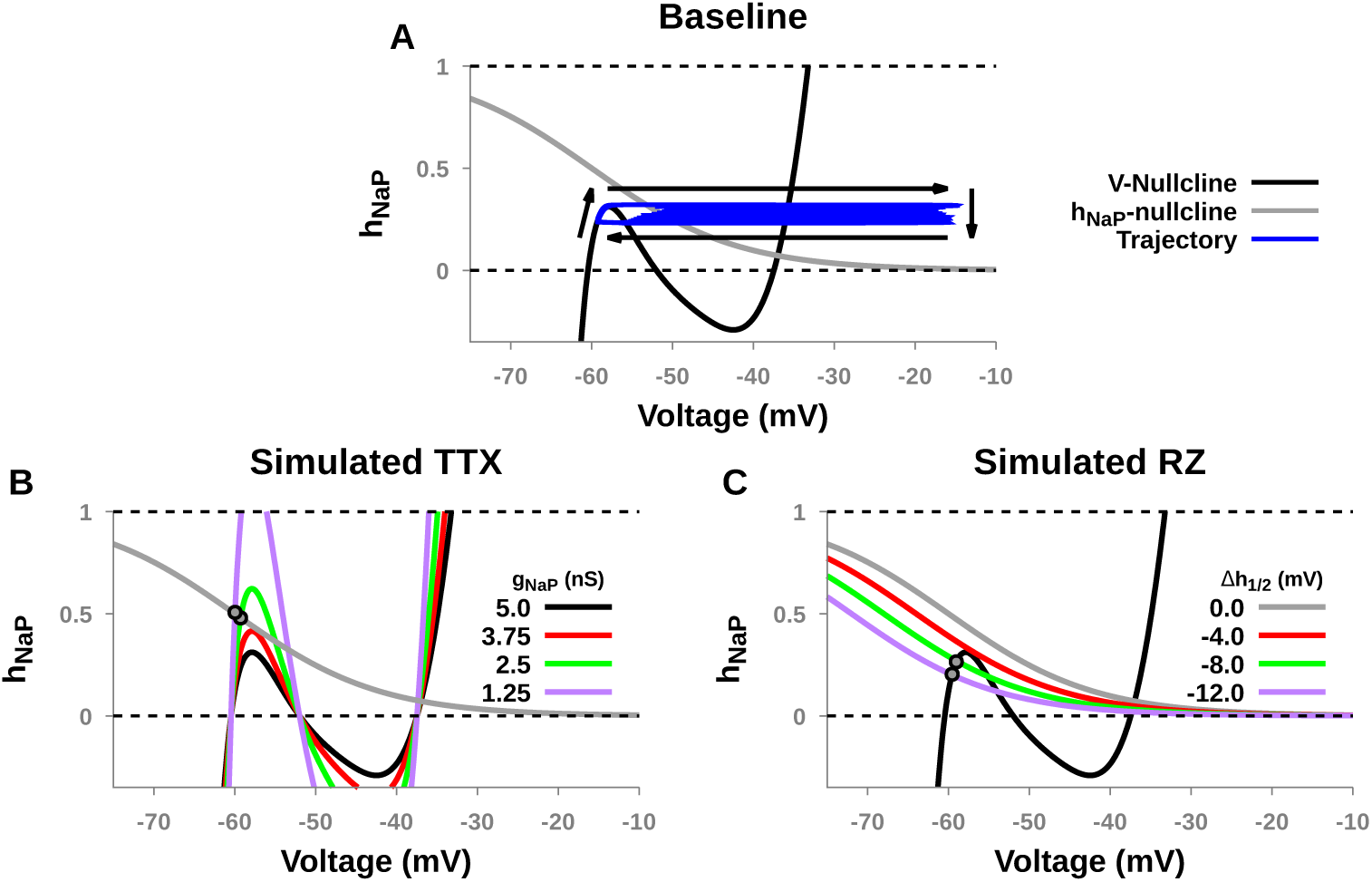
Phase plane representation of simulated TTX and RZ block of *I*_*NaP*_ in an isolated pre-BötC neuron. (A) *V*_*m*_- and *h*_*NaP*_ -nullclines (black and gray, respectively) projected to the (*V*_*m*_, *h*_*NaP*_)-plane under control conditions (*g*_*NaP*_ = 5.0 *nS*) in a bursting regime, along with a projection of the bursting trajectory (blue). Note that this view does not represent the *I*_*NaP*_ activation variable *m*_*NaP*_. (B) Increasing TTX blockade of *I*_*NaP*_ moves the left knee of the *V*_*m*_-nullcline to larger values of *h*_*NaP*_, which also abolishes bursting by creating a stable fixed point on the left branch of the *V*_*m*_-nullcline. (C) Increasing RZ blockade of *I*_*NaP*_ shifts the h-nullcline to the left, abolishing intrinsic bursting by creating a stable fixed point left branch of the *V*_*m*_-nullcline. *g*_*TonicE*_ = 0.35 *nS* in all panels.

### Transient hyperpolarization elicits rebound bursting after *I*_*NaP*_ blockade

Although both forms of *I*_*NaP*_ blockade convert the model neuron’s intrinsic dynamics from bursting to silence, we next considered whether transient bursting could be elicited by perturbations. As can be seen in Fig 4A, a burst occurs when *h*_*NaP*_ rises to a value above that of the left knee of the *V*_*m*_-nullcline, call it 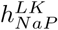, such that the model’s trajectory can transition to high voltages. The position of the entire *V*_*m*_-nullcline, including its left knee, depends not only on the extent of *I*_*NaP*_ blockade but also on the level of input to the neuron. Indeed, hyperpolarizing inputs shift the *V*_*m*_-nullcline to more hyperpolarized *V*_*m*_ values. As a result, its left branch equilibrium point ends up more hyperpolarized in *V*_*m*_ and, due to the voltage-dependence of *I*_*NaP*_, at a more deinactivated, larger *h*_*NaP*_ value, call it 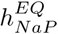. The left knee of the *V*_*m*_-nullcline also rises to a new value, 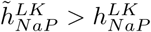. In theory, as the model settles toward the equilibrium 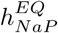, although *h*_*NaP*_ remains below 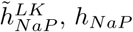 could exceed the original left knee height, 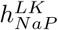. If this happens, then release from hyperpolarization will result in a single transient rebound burst. For this result to occur, three conditions need to be met: (1) 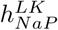 must be less than 1, so that it is physically realizable, (2) the hyperpolarization must be sufficiently strong that 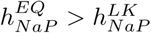, and (3) the hyperpolarization must be maintained long enough.

Fig 5 characterizes the magnitude and duration of hyperpolarization needed to elicit rebound bursting as a function of simulated TTX and RZ blockade of *I*_*NaP*_ in our model. We found that with simulated TTX blockade of *I*_*NaP*_ rebound bursting is still possible over narrow range of *g*_*NaP*_, *g*_*NaP*_*∈* (1.6*-*3.55 *nS*). The magnitude and duration of hyperpolarizing input needed to elicit rebound bursting quickly becomes biologically unrealistic, however. In contrast, there are always input magnitudes and durations that yield rebound bursting following any level of simulated RZ blockade, because RZ blockade does not affect 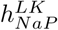 and thus condition (1), 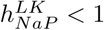, holds for all levels of Δ*h*_1*/*2_.

**Fig 5.**
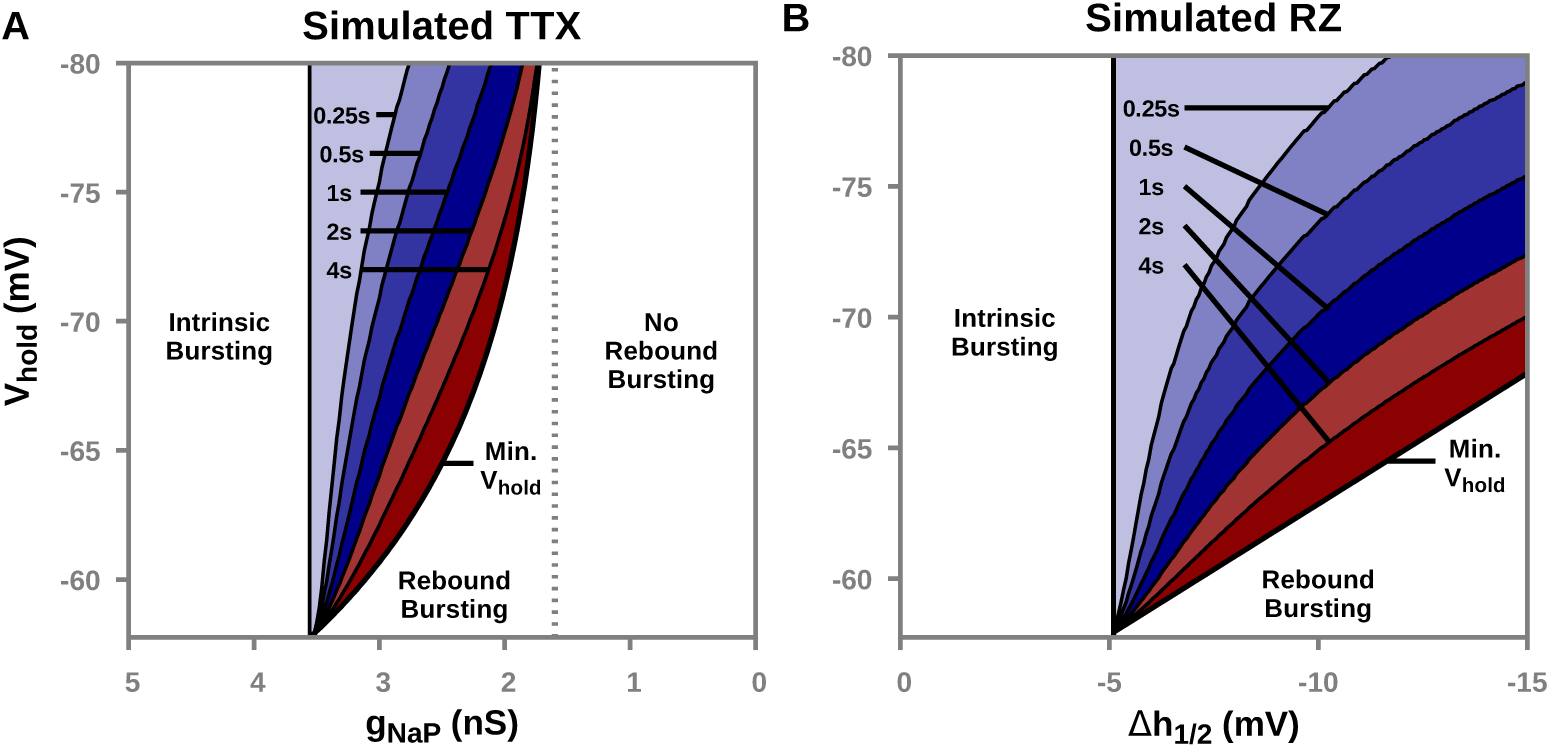
Hyperpolarized holding potential (*V*_*hold*_) and duration (color-coded) required to elicit rebound bursting as a function of simulated (A) TTX or (B) RZ blockade of *I*_*NaP*_. Here, *g*_*Tonic*_ = 0.35 *nS*.

In in vivo conditions, the pre-BötC receives strong inhibition during the interburst interval from other nuclei involved in respiratory rhythm and pattern formation [13, 17, 18, 19, 20]. During this interval the transient hyperpolarization is approximately 10 *mV* in magnitude and 2 *s* in duration. Our simulations predict that ”complete” blockade of *I*_*NaP*_, defined by complete loss of sustained intrinsic bursting (Fig 3), will stop this inhibitory input from inducing a burst in pre-BötC neurons if the block is applied via TTX (Fig 6). This loss occurs because after TTX application, the hyperpolarization does not achieve condition (2), 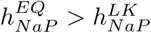. If the block is induced with RZ, however, then the trajectory reaches a position in the (*V*_*m*_, *h*_*NaP*_) plane where condition (2) does hold, such that subsequent removal of hyperpolarization will result in a burst. Thus, these simulations predict that in the presence of repetitive cycles of inhibition associated with ongoing respiratory rhythms in vivo, pre-BötC neurons will still generate bursts supported by *I*_*NaP*_ even after RZ application.

**Fig 6.**
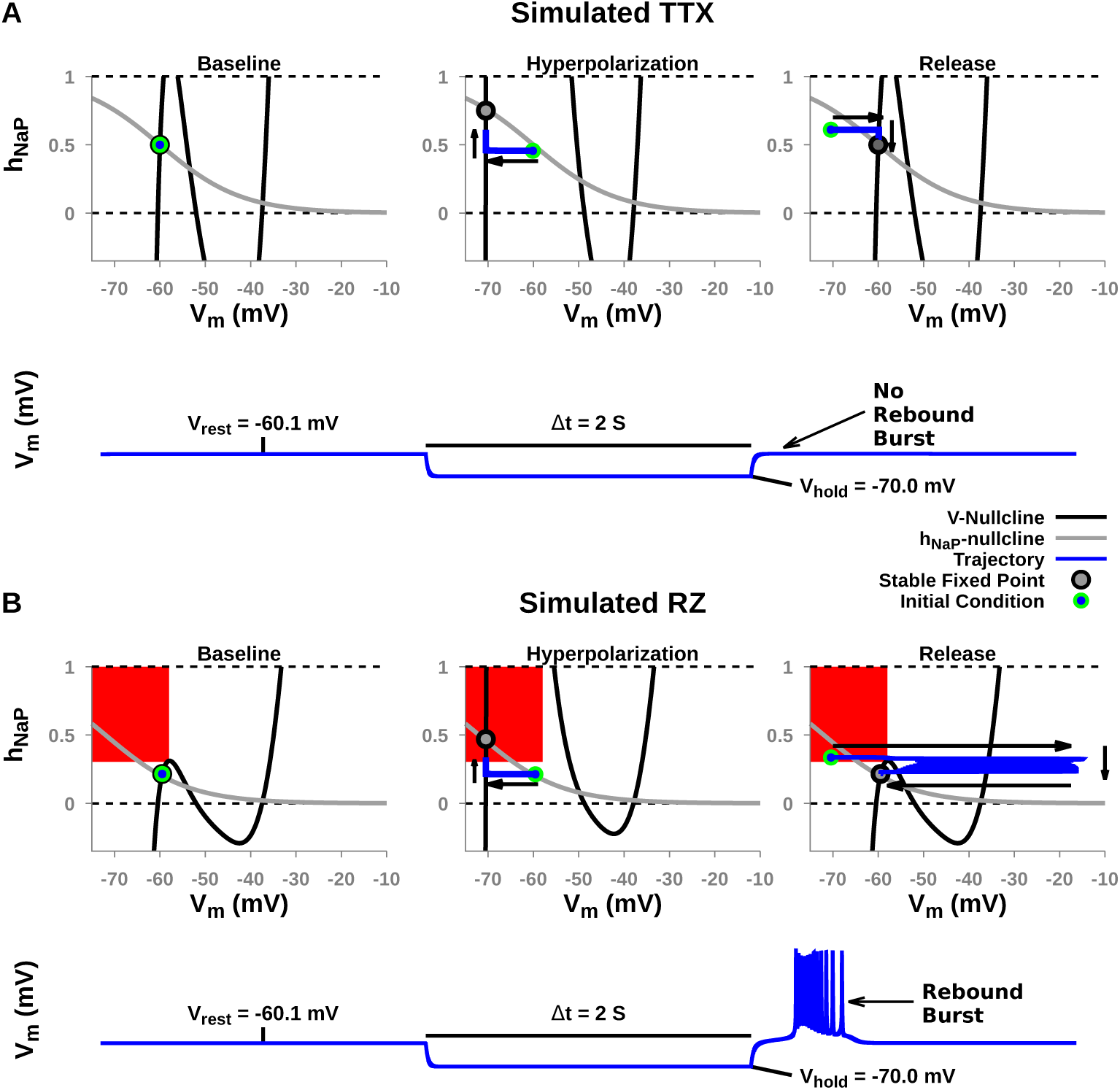
A transient hyperpolarizing perturbation fails to elicit rebound bursting after simulated (A) TTX but not (B) RZ blockade of *I*_*NaP*_. (A & B) Top Panels: *V*_*m*_- and *h*_*NaP*_ -nullclines during baseline, hyperpolarization, and release from hyperpolarization. Arrows indicate the direction of the trajectory. (A & B) Bottom Panels: voltage trace of *V*_*m*_ during baseline hyperpolarization and release. Initial conditions that generate transient bursting trajectories are indicated by the filled red regions and only appear in the RZ case. For simulated TTX blockade in (A) *g*_*NaP*_ = 1.25 *nS*. In this case, after hyperpolarization (middle panel), the equilibrium point (open circle) lies at an *h*_*NaP*_ value that is too low to allow clearance of the left knee after release (right panel). For simulated RZ blockade in (B), Δ*h*_1*/*2_ = *-*12 *mV*. Here, 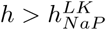 in the red region, and hence any hyperpolarization that allows the trajectory to enter the red region (e.g., center panel) will yield a burst upon release (right panel). For both simulations, *g*_*TonicE*_ = 0.35 *nS*.

### The pre-BötC pre-I network

Next, to investigate simulated TTX and RZ application in the pre-BötC network, we constructed a heterogeneous population of 50 model pre-BötC neurons coupled though all-to-all excitatory synapses; see *Materials and Methods* for a full model description. This network is often referred to as the pre-inspiratory/inspiratory (pre-I/I) population and is thought to drive the fictive inspiratory rhythm seen in in vitro slice preparations [13, 22, 23]. For consistency with experimental observations, parameters were chosen such that approximately 30% of neurons in the synaptically uncoupled network remain rhythmically active [24]; for simplicity, we also tuned the synaptically coupled network such that all neurons are recruited into network oscillations (Fig 7).

**Fig 7.**
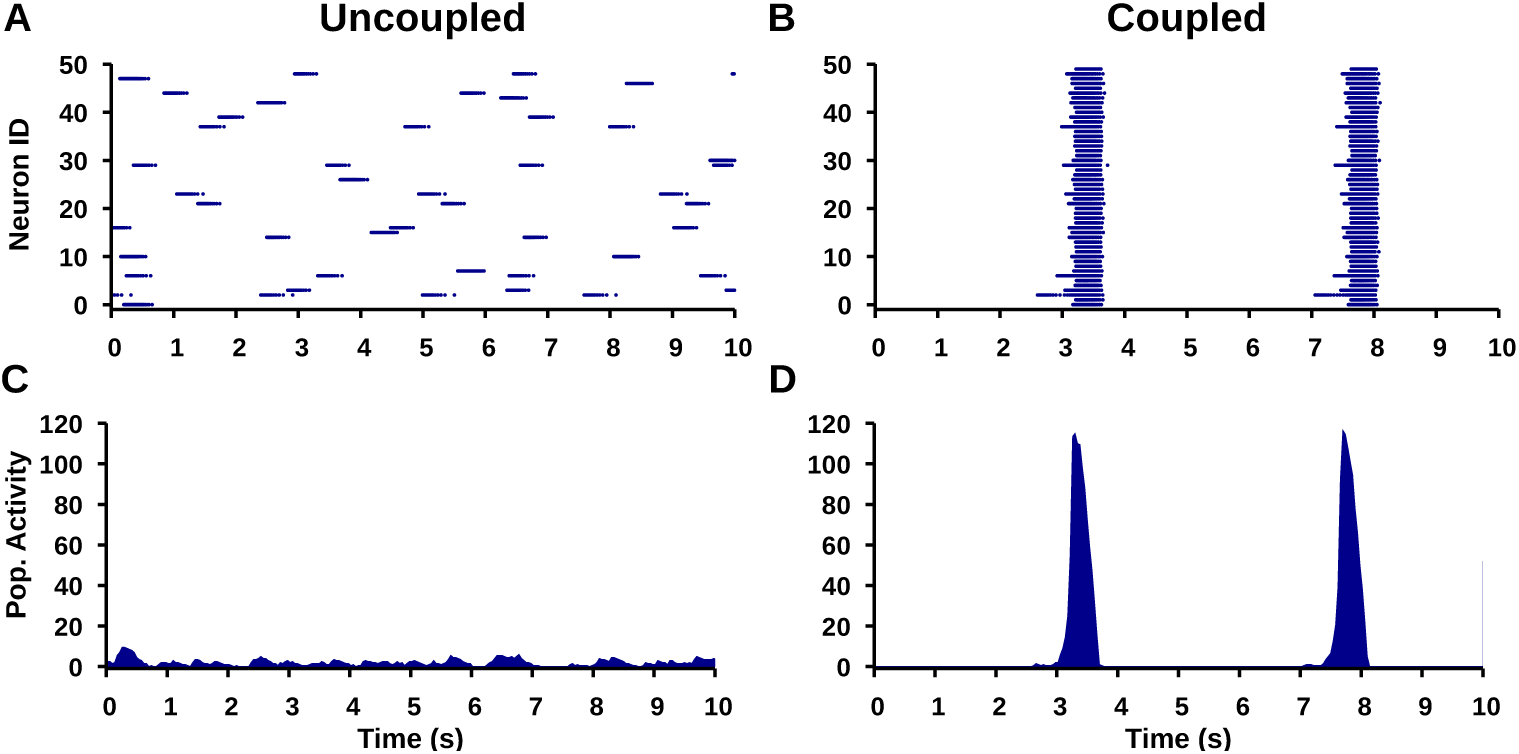
Simulated pre-BötC network activity. (top) Raster plots showing (A) intrinsic bursting in a subset of neurons in the synaptically uncoupled network and (B) synchronized network bursts in the synaptically coupled network. (bottom) Integrated population activity for the (C) uncoupled and (D) coupled network.

### Uniform TTX and RZ block of *I*_*NaP*_ in the pre-BötC pre-I network

The synaptic coupling between neurons within the pre-BötC network does not alter the mechanisms of action of TTX and RZ on voltage-gated sodium channels described above. We found that increasing the degree of uniform *I*_*NaP*_ block by simulated TTX or RZ results in a progressive reduction in network frequency and a slight reduction in network amplitude, followed by an abrupt cessation of network oscillations (Fig 8 A & B). With RZ, however, additional off-target effects need to be considered [10, 25]. Specifically, at the same concentrations (0*-*20*µM*) used to block *I*_*NaP*_ in respiratory circuits [9, 12, 13], RZ also inhibits glutamatergic excitatory synaptic transmission [11, 26]. To understand this off-target effect, *I*_*SynE*_ was blocked in a separate set of simulations by systematically reducing the excitatory synaptic conductance parameter (*W*_*MaxE*_). In contrast to the *I*_*NaP*_ effects, we found that simulating the progressive block of synaptic excitation by RZ results in a slight increase in network frequency and a large reduction in network amplitude (Fig 8 C). These outcomes agree with the results from experimental block of excitatory synapses in the pre-BötC [23].

**Fig 8.**
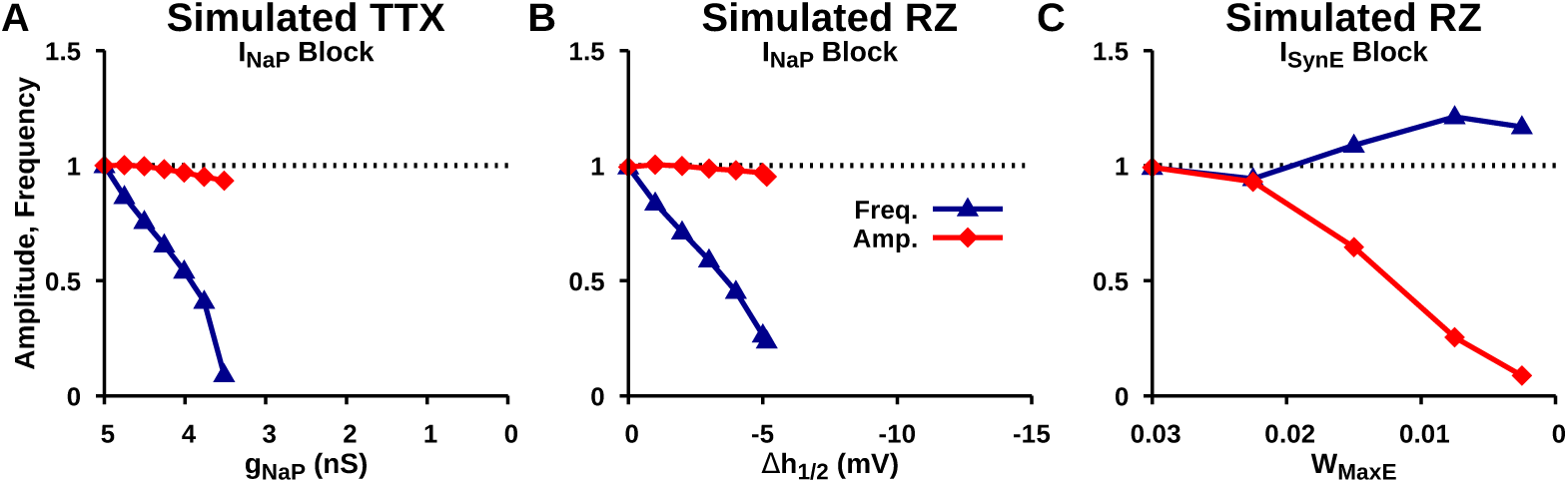
Effects of simulated (A) TTX and (B) RZ block of *I*_*NaP*_ as well as (C) RZ block of *I*_*SynE*_ on network amplitude and frequency of network oscillations in the pre-BötC pre-I neuron population.

Next, we compared simulated uniform TTX and RZ application with experimental data adapted from [9]. This experimental dataset characterizes the dose-dependent effects of bilateral microinfusion of TTX and RZ within the pre-BötC of neonatal rat brainstem slices in vitro on the amplitude and frequency of ∫XII motor output as a function time. For the comparison of experimental and simulated data, we plotted amplitude versus frequency in order to eliminate temporal dynamics. Consistent with the experimental data, we found that simulated TTX application in the pre-BötC progressively reduces the frequency without affecting the amplitude of population oscillations before oscillations abruptly stop (Fig 9 A). With RZ, however, assuming that both *I*_*NaP*_ and excitatory synapses are impacted, extrapolation from our separate simulations of these effects predicts that experimental RZ application will progressively reduce both frequency and amplitude of the pre-BötC population oscillations until they eventually stop. Fig 9 B demonstrates that these effects do arise both in simulated RZ application, where both *I*_*NaP*_ and synapses are affected, and in experimental data [9]. We found that our simulation results matched the reduction in network amplitude and frequency seen with experimental RZ application when *W*_*MaxE*_ decayed exponentially with increasing hyperpolarization of 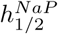 (see *Materials and Methods*). To match experimental results of RZ application at 5 *µM, W*_*MaxE*_ was maximally reduced by 7% (Fig 9 B, Tuning 1), and for RZ at both 10 *µM* and 20 *µM, W*_*MaxE*_ was maximally reduced by 15% (Fig 9 B, Tuning 2).

To investigate the mechanisms underlying changes in network amplitude and frequency, we quantified changes in the *I*_*NaP*_ threshold 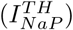 required required for spike generation and the peak *I*_*SynE*_ magnitude during network bursts, and we estimated the *h*_*NaP*_ threshold 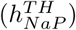 required to initiate bursting as a function of the level of TTX and RZ application in our simulations, tuned to match experimental data from [9] (Fig 9 a1, b1, b2).

**Fig 9.**
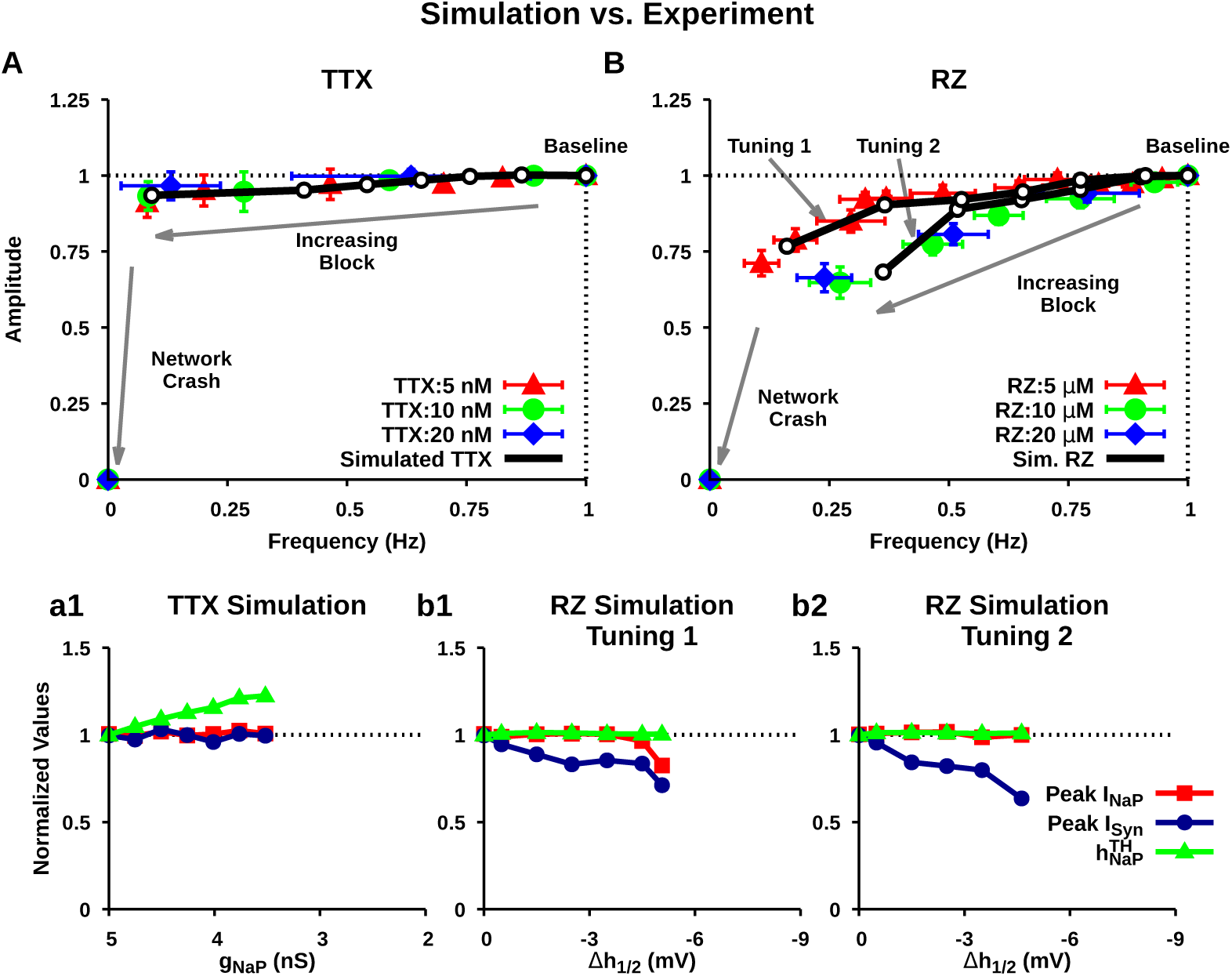
Comparison of experimental (colored) and simulated (black) effects of (A) TTX and (B) RZ application in the pre-BötC on the amplitude and frequency of pre-I network oscillations. Simulated RZ application affects *I*_*NaP*_ and excitatory synapses whereas TTX only affects *I*_*NaP*_. In the simulations, the relevant values at the end points (where network oscillations stop) are as follows: (A) *g*_*NaP*_ = 3.52; (B) (top trace) Δ*h*_1*/*2_ = *-*5.0 *mV, W*_*maxE*_ = 0.0255 *nS*, (bottom trace) Δ*h*_1*/*2_ = *-*4.5 *mV, W*_*maxE*_ = 0.0224 *nS*. Experimental data was adapted from [9] and shows the progressive change in amplitude and frequency of network oscillations (monitored by integrated hypoglossal nerve activity) relative to baseline following bilateral microinfusion of TTX or RZ at different concentrations into the pre-BötC. Notice that, TTX only effects frequency, whereas RZ affects frequency and amplitude. (a1, b1, b2) Effect of simulated TTX and RZ (tunings 1 & 2) blockade on the peak *I*_*NaP*_, peak *I*_*Syn*_, and the *I*_*NaP*_ inactivation threshold 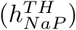 required to initiate bursting. 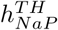 was defined as the maximal value of the mean population *h*_*NaP*_ prior to burst initiation.

In the model, amplitude is defined as the number of spikes per neuron per 50 *ms* bin. Therefore, by this definition, changes in network amplitude result from changes in the firing rate of bursting neurons and/or changes in the number of neurons recruited into network bursts. The number of recruited neurons is affected by *I*_*SynE*_ and the firing rate of individual neurons during bursting is a function of the total depolarizing current, determined by both *I*_*NaP*_ and *I*_*SynE*_. In contrast, network oscillation frequency is determined by the time required for *I*_*NaP*_ to recover from inactivation to a sufficient threshold 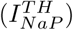 for spike generation following each network burst. To be more precise, network frequency is a function of the time required for *h*_*NaP*_ to recover from inactivation up to a level, 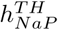, such that 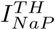 is reached following each network burst. For a fixed level of synaptic input, threshold 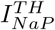 is a constant given by 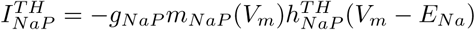 (see equation 5 in *Materials and Methods*).

Thus, as long as synaptic input remains constant, 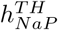 increases as *g*_*NaP*_ decreases. In the period of *h*_*NaP*_ recovery leading up to burst onset, we indeed have the constant synaptic input *I*_*SynE*_ = 0, independent of *W*_*MaxE*_, since the pre-BötC neurons activate synchronously in the regime we are considering.

In the case of simulated TTX, while 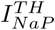 and *I*_*SynE*_ did not change, the above reasoning yields an increase in 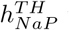 with the progressive decrease in *g*_*NaP*_ (Fig 9 a1). Note that the increase in the 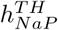 is consistent with Fig4. Therefore, the reduced network frequency seen with increasing TTX blockade is a direct consequence of an increased *h*_*NaP*_ recovery time driven by the increased 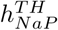. Because 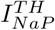 does not vary with *g*_*NaP*_, the level of *I*_*NaP*_ expressed in bursting neurons is the same as before the blockade. This invariance of *I*_*NaP*_ together with the fact that TTX does not affect *I*_*syn*_ imply that the firing rate of bursting neurons and the number of recruited neurons are unchanged (not shown). This reasoning explains why network amplitude is unaffected by simulated TTX blockade of *I*_*NaP*_.

In the case of simulated RZ, since *g*_*NaP*_ does not vary, 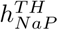 is unaffected by progressive blockade. In this scenario, changes in frequency arise instead due to a slower rate of recovery from inactivation caused by the hyperpolarizing shift in the inactivation curve (i.e., the *h*_*NaP*_ -nullcline), Fig 2 & 4. This shift is accompanied by a reduction of *I*_*SynE*_ that follows directly from the reduction of *g*_*SynE*_ with simulated RZ. Therefore, changes in network amplitude with simulated RZ arise due to the progressive reduction of *I*_*SynE*_, which results in a decreased firing rate during bursting and de-recruitment of a subset of the neurons (data not shown).

### Non-uniform TTX and RZ block of *I*_*NaP*_ in the pre-BötC pre-I network

Blockade of *I*_*NaP*_ by experimental application of TTX or RZ is likely to be non-uniform, especially in the case of in vitro bath application where drug penetration depends on passive diffusion. With TTX and RZ bath application, *I*_*NaP*_ will be blocked in neurons closest to the surface before neurons at the center of the slice. As mentioned previously, the pre-BötC pre-I population contains pacemaker and non-pacemaker neurons that play different roles in rhythm and pattern formation. The spatial orientation of these two types of neurons within the pre-BötC is unknown. Therefore, to understand the effects of non-uniform drug penetration in this model, we simulated the sequential blockade of *I*_*NaP*_ by TTX and RZ in the pre-I population, under each of three different assumptions on the order in which pacemaker and non-pacemaker neurons are affected: (1) pacemaker then non-pacemaker, (2) non-pacemaker then pacemaker, and (3) random order. We found that the effects of TTX and RZ are indistinguishable for all three cases. If pacemaker neurons are affected first, then *I*_*NaP*_ block via either mechanism results in a progressive reduction in network frequency with no effect on amplitude until oscillations eventually stop (Fig 10A). That is, with loss of *I*_*NaP*_ in pacemakers, burst initiation is slowed, but once a burst occurs, all neurons are recruited. In contrast, if non-pacemaker neurons are affected first, then *I*_*NaP*_ block results in a progressive reduction in network amplitude and an increase in frequency before oscillations eventually stop (Fig 10B). The loss of *I*_*NaP*_ in non-pacemakers can prevent their recruitment, while the involvement of fewer neurons in each burst results in shorter bursts that can recur more frequently. Finally, if *I*_*NaP*_ is blocked in pacemaker and non-pacemaker neurons in a random order, then *I*_*NaP*_ block results in a progressive reduction in network amplitude and frequency before oscillations eventually stop. In all cases, network oscillations stop as soon as *I*_*NaP*_ is completely blocked in all pacemaker neurons.

**Fig 10.**
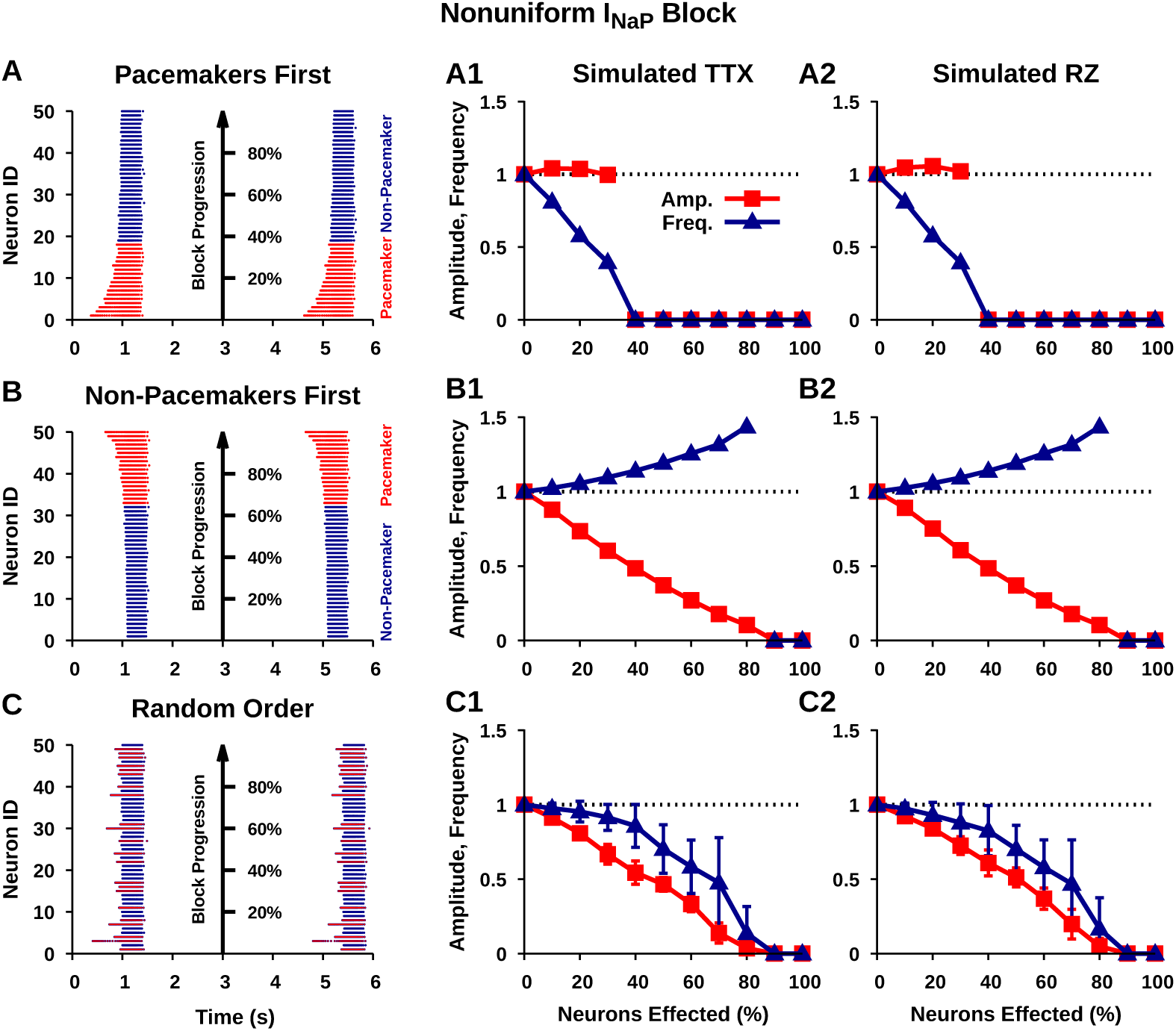
Simulated progressive nonuniform *I*_*NaP*_ block in the isolated pre-I network. (A,B,C) From left to right: network burst with neurons color-coded as pacemakers (red) or non-pacemakers (blue) and numbered based on order affected from 1 (first affected) to 50 (last affected); amplitude and frequency of isolated pre-I population oscillations as a function of the percentage of the pre-I population where *I*_*NaP*_ is completely blocked. For a given neuron, *I*_*NaP*_ is considered completely blocked when *g*_*NaP*_ = 0 *nS* in the case of TTX (A1,B1,C1) or when Δ*h*_1*/*2_ = *-*15 *mV* and *I*_*SynE*_ is attenuated by 25% (A2,B2,C2), see Figs. 2, 8. Error bars in C1 and C2 indicate the *±*SEM of ten trials where *I*_*NaP*_ is progressively blocked across the network in random order.

### Simulated TTX and RZ block of *I*_*NAP*_ in the intact respiratory network

Finally, to investigate the effects of simulated TTX and RZ block of *I*_*NaP*_ in the intact respiratory network, we reconstructed a multi-population network model that is thought to represent the core mammalian respiratory central pattern generator located in the BötC and pre-BötC [13, 17, 27, 28, 29, 30]. The simulated intact network is composed of four subpopulations each consisting of 50 neurons. The subpopulations are characterized by the timing of their activity relative to the phases of inspiration and expiration as follows: post inspiratory (post-I), augmenting expiratory (aug-E), pre-I/I as simulated in the pre-BötC network considered thus far in this work, and early-inspiratory (early-I). This network produces a three-phase rhythm with inspiratory (I), post-inspiratory (pI), and stage-2 expiratory (E2) phases, which is similar to respiratory rhythms seen in vivo (Fig 11). For a full model description, see *Materials and Methods*.

**Fig 11.**
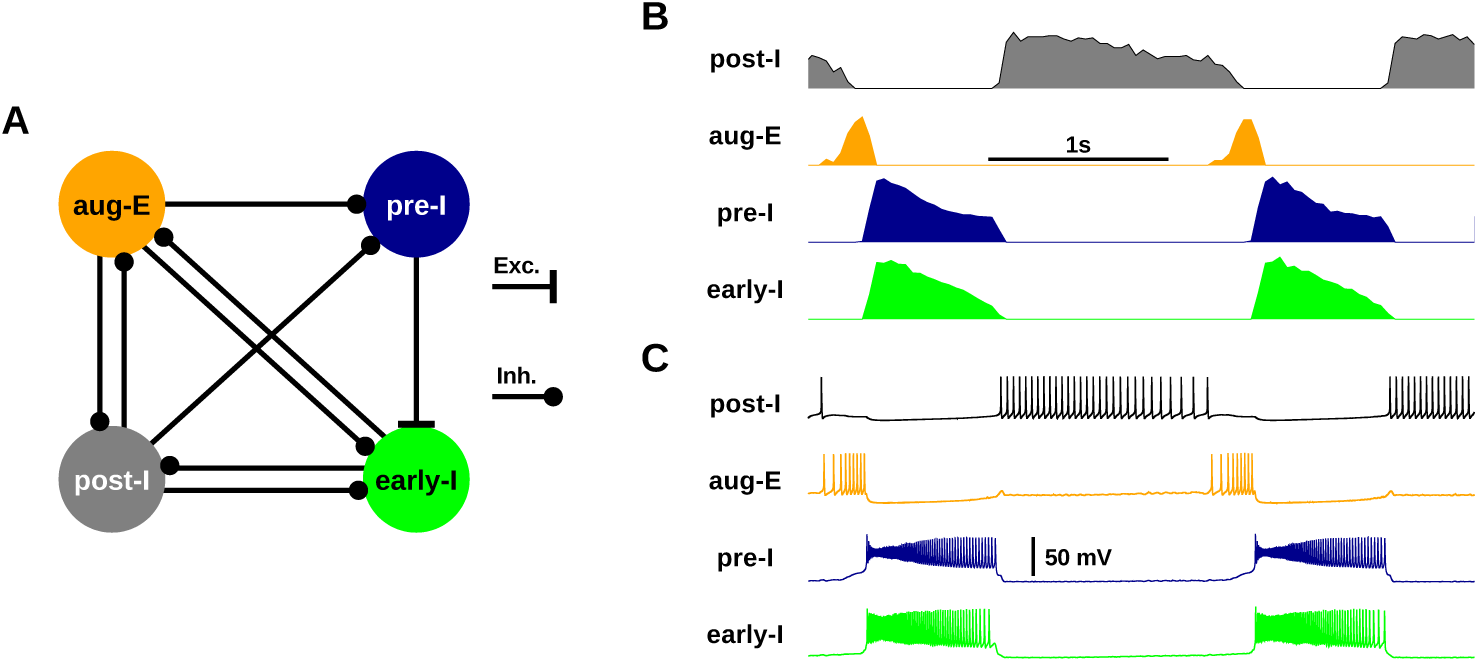
Simulated intact respiratory network. (A) Circuit diagram of the intact respiratory network composed of the inhibitory (post-I, aug-E early-I) and excitatory (pre-I) subpopulations. (B) Integrated subpopulation spiking activity. Amplitudes of each subpopulation were normalized. (C) Spiking in example neurons from each subpopulation.

In the intact respiratory network, the respective contributions of *I*_*NaP*_ and inhibitory network interactions to rhythm generation vary depending on the excitability state of the pre-I population [28]. Therefore, before simulating TTX and RZ blockade of *I*_*NaP*_ in the intact network, we first characterized the dependence of pre-I population bursting on *I*_*NaP*_ and synaptic inhibition in the intact network as a function of *g*_*Tonic*_ (Fig 12 A). This was accomplished by comparing the dynamics of the pre-I population (silent, bursting, tonic), embedded in the network, between baseline conditions and the extreme cases where *I*_*NaP*_ = 0 or *I*_*SynE*_ = 0. We found that under baseline conditions, bursting in the pre-I population is extremely robust and oscillations continue over all tested values of *g*_*Tonic*_, namely 0.3*-*1.0 *nS* (cf. [31]). Complete block of inhibitory synaptic currents reveals that the pre-I population bursting is dependent on inhibitory network interactions for *g*_*Tonic*_ *∈* (0.40, 1), indicated by a transition of the pre-I population from bursting, under baseline conditions, to a tonic spiking mode of activity when *I*_*SynE*_ = 0. In contrast, complete blockade of *I*_*NaP*_ reveals a regime of *I*_*NaP*_ -dependent bursting for *g*_*Tonic*_ = 0.3 *-* 0.486, indicated by a transition of the pre-I population from a bursting to a silent mode of activity when *I*_*NaP*_ = 0. Importantly, these two regimes overlap (for *g*_*Tonic*_ *∈* (0.40, 0.486)), which indicates a region where both *I*_*NaP*_ and inhibitory synaptic interactions are critical for rhythm generation/bursting in the pre-I population (Fig 12 A).

**Fig 12.**
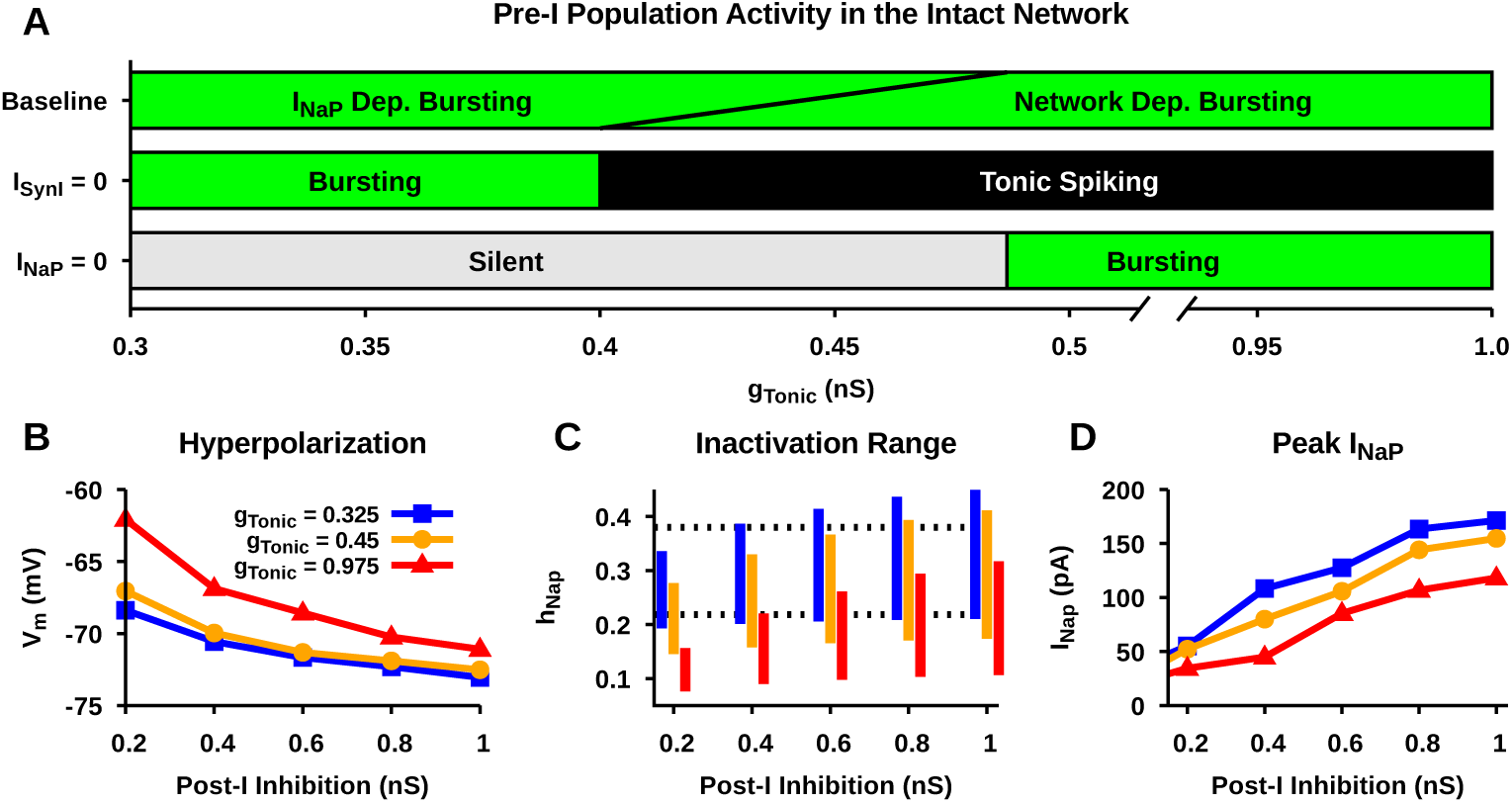
Characterization of the intact respiratory network behavior. (A) Identification of *I*_*NaP*_ and/or network dependent regimes of pre-I population bursting as a function of tonic excitatory drive (*g*_*Tonic*_). Effect of post-I inhibition strength and *g*_*Tonic*_ on (B) post-inspiratory phase hyperpolarization, (C) *h*_*NaP*_ dynamic range, and (D) the peak inspiratory phase *I*_*NaP*_ in the pre-I population. Notice that the *I*_*NaP*_ level is strongly affected by the magnitude of post-I inhibition and *g*_*Tonic*_. Strong post-I inhibition increases the magnitude of post-inspiratory phase hyperpolarization, which decreases *I*_*NaP*_ inactivation and increases the peak *I*_*NaP*_. In contrast, strong *g*_*Tonic*_ decreases post-inspiratory phase hyperpolarization, which increases *I*_*NaP*_ inactivation and decreases the peak *I*_*NaP*_. Dashed lines in C indicate the dynamic range of *h*_*NaP*_ in the isolated pre-I population under baseline conditions used in Fig 7.

Next we characterized network parameters affecting the magnitude/contribution of *I*_*NaP*_ during pre-I population bursting, which, as in the pre-I network, modulate *I*_*NaP*_ inactivation dynamics. As with the pre-I network, there is a threshold level of *I*_*NaP*_ needed for pre-BötC burst onset, which is realized when *h*_*NaP*_ deinactivates to a corresponding threshold level, 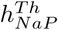. One new feature in this case, however, is that 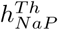 depends both on *g*_*Tonic*_ and on the level of synaptic inhibition to the pre-BötC population, since this parameter and input affect the position of the *V*_*m*_-nullcline for the pre-BötC neuron (analogously to the impact of applied inhibitory current on a single pre-BötC neuron shown in Fig 6B). This dependence holds in both the *I*_*NaP*_ -dependent and the network-dependent burst regimes (Fig 12). Therefore, in the pre-I population, we characterized: (1) the maximal hyperpolarization during the fictive expiratory phase, which impacts the rate of deinactivation of *h*_*NaP*_, (2) the dynamic range of *h*_*NaP*_ over a complete respiratory cycle, and (3) the peak *I*_*NaP*_ during bursting, as a function of the strength of inhibition from the post-I population for three levels of pre-I population tonic excitation set by *g*_*Tonic*_ (Fig 12B, C and D).

Increasing *g*_*Tonic*_ naturally decreases the hyperpolarization of the pre-I neurons in the expiratory phase (Fig 12B). Due to its effects on the *V*_*m*_-nullclines for the pre-I neurons, it also lowers 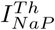 and hence 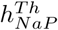. These changes lower the operating range of *h*_*NaP*_ (Fig 12C) and correspondingly result in a lower level of *I*_*NaP*_ while the pre-I population is active (Fig 12D). In contrast, increasing the strength of inhibition from the post-I population to the pre-I neurons hyperpolarizes them more (Fig 12B) and increases 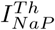 and hence 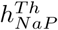 through the effect of inhibition on the pre-I *V*_*m*_-nullcline. These changes raise the operating range of *h*_*NaP*_ to higher levels (Fig 12C) and correspondingly allow *I*_*NaP*_ to reach a higher level. Note that the contribution of *I*_*NaP*_ is small when *g*_*Tonic*_ is relatively strong and post-I inhibition is weak, whereas the contribution of *I*_*NaP*_ is large when the relative strengths of *g*_*Tonic*_ and post-I inhibition are reversed. Interestingly, under the latter conditions, the magnitude of *I*_*NaP*_ can be larger in the intact network than in the isolated pre-I population due to the post-I population inhibition, which can enhance *I*_*NaP*_ deinactivation.

Finally, we simulated TTX and RZ blockade of *I*_*NaP*_ in the intact respiratory network (Fig 13). The effects of simulated *I*_*NaP*_ blockade on the pre-I population dynamics are expected to depend on the value of *g*_*Tonic*_ and the strength of post-I inhibition within this population, due to the effects illustrated in Fig 12. Therefore, we characterized the relative change in the pre-I population amplitude and frequency as a function of *g*_*Tonic*_ and post-I inhibition for ’complete’ blockade of *I*_*NaP*_ by simulated TTX and RZ. Blockade of *I*_*NaP*_ was considered complete for simulated TTX when *g*_*NaP*_ = 0 *nS*, and for simulated RZ when Δ*h*_1*/*2_ = *-*15 *mV* (see Fig 2).

**Fig 13.**
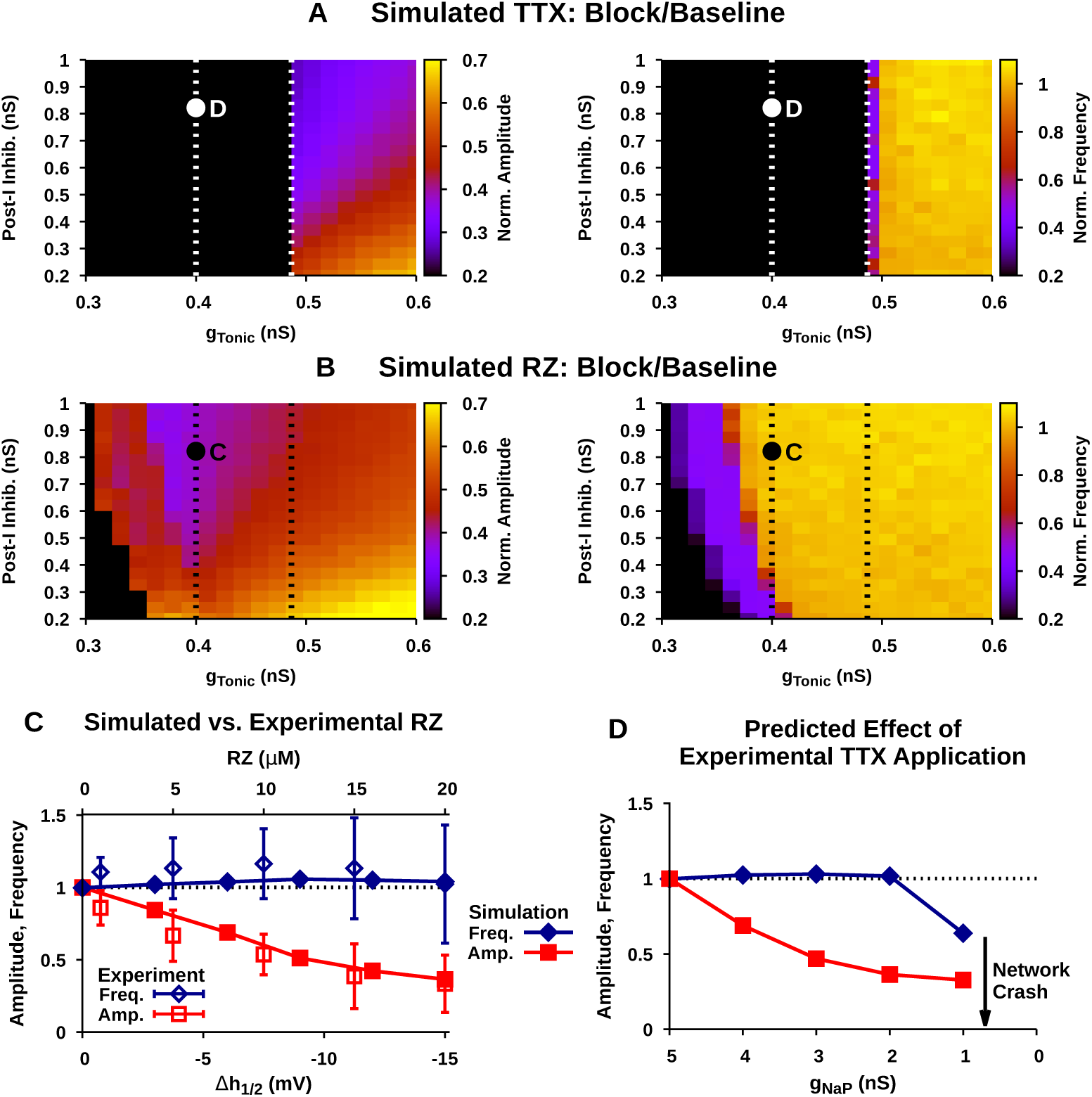
Predicted effect of blockade of *I*_*NaP*_ in the intact network. Relative effects of ”complete” blockade of *I*_*NaP*_ by simulated (A) TTX and (B) RZ application on the amplitude (left) and frequency (right) as a function of tonic excitatory drive (*g*_*Tonic*_) and the strength of post-I inhibition. Oval shape indicates parameters capable of matching the maximal relative changes in amplitude and frequency seen with experimental application of RZ. The points labeled C and D indicate the *g*_*Tonic*_ and post-I inhibition values used in panels C and D. (C) Comparison of experimental and simulated RZ blockade of *I*_*NaP*_. Simulated RZ application closely matches experimental data when *g*_*Tonic*_ = 0.425 and post-I inhibition = 0.8. Experimental data is adapted from [13]. (D) Predicted effect of experimental TTX blockade of *I*_*NaP*_ in the intact network under identical conditions to panel C.

In the case of simulated TTX, complete blockade of *I*_*NaP*_ abolishes pre-I population oscillations for *g*_*Tonic*_ *<* 0.486 *nS* across all levels of post-I inhibition considered (Fig 13A), as expected from the boundary between *I*_*NaP*_ - and network-dependent bursting regions in Fig 12A. In contrast, for *g*_*Tonic*_ *>* 0.486 *nS* pre-I population oscillations continue after complete blockade of *I*_*NaP*_. In this regime, network amplitude is generally reduced without affecting frequency, except for a frequency reduction for *g*_*Tonic*_ ≈ 0.486 *nS*. The extent of the reduction in amplitude depends *g*_*Tonic*_ and the strength of post-I inhibition and is largest when *g*_*Tonic*_ is weak and post-I inhibition is strong.

In the case of simulated RZ, the range of *g*_*Tonic*_ values where complete blockade of *I*_*NaP*_ abolishes pre-I population bursting is greatly reduced (Fig 13B) compared to simulated TTX. In general, stronger post-I inhibition decreases the range of *g*_*Tonic*_ values where *I*_*NaP*_ block abolishes pre-I population bursting. This relationship is due to the increased recovery of *I*_*NaP*_ from inactivation caused by the strengthening of post-I inhibition and the enhanced expiratory phase hyperpolarization of pre-I that it induces. This mechanism is illustrated in Fig 6. In the region of parameter space where pre-I population bursting is not abolished, network amplitude is generally reduced by RZ blockade without affecting frequency. An exception occurs close to the boundary where bursting is almost abolished; in this case RZ blockade reduces both amplitude and frequency. Moreover, the relative decrease in amplitude is largest when *g*_*Tonic*_≈0.4 *nS* and post-I inhibition is strong (≿0.5 *nS*), Fig 13. Importantly, in these simulations, complete blockade of *I*_*NaP*_ by RZ fails to stop the respiratory rhythm under network conditions where pre-I population bursting is dependent on *I*_*NaP*_ (Fig 12A).

Finally, we compared the effects of simulated RZ blockade of *I*_*NaP*_ in the intact respiratory network with experimental data from [13] (Fig 13C). This experimental data set characterizes the steady-state dose-dependent effects of RZ on burst amplitude and frequency of the intact respiratory network output measured from integrated phrenic nerve activity. In these experiments, the maximal 20 *µm* RZ concentration, which is assumed to completely block *I*_*NaP*_, resulted in a *∼*35% reduction in network amplitude and no significant change in frequency. In order to match these data in our simulations, we found that post-I inhibition must be relatively strong (*≳*0 5 *nS*) and *g*_*Tonic*_ between 0.375 *nS* and 0.425 *nS* (Fig 13B, C). Interestingly, this range of *g*_*Tonic*_ values overlaps with the regimes where pre-I population bursting is exclusively *I*_*NaP*_ -dependent and where bursting is dependent on both *I*_*NaP*_ and inhibitory network interactions (Fig 12A). Consequently, under these network conditions, our model predicts that selective and complete block of *I*_*NaP*_ by experimental application of TTX at low concentrations (*≤*20 *nM*) will abolish the respiratory rhythm in the intact network (Fig 13D).

## Discussion

Understanding how pharmacological mechanisms and network dynamics affect *I*_*NaP*_ blockade by TTX and RZ is critical for interpreting experimental data and its implications for understanding the underlying mechanisms of respiratory rhythm and pattern formation. Therefore, in this computational study, we characterized the effects of TTX and RZ blockade of *I*_*NaP*_ on respiratory network dynamics by simulating their distinct pharmacological mechanisms of action in established models of respiratory neurons and neurocircuitry that represent in vitro and in vivo mouse/rat preparations. To summarize, we show that in simulated pre-BötC respiratory neurons under conditions representing in vitro slice preparations, TTX and RZ both effectively block *I*_*NaP*_ and abolish intrinsic bursting (Figs 2 & 3). Given these findings, it is tempting to conclude that TTX and RZ application will induce similar effects on respiratory dynamics under all experimental conditions. Interestingly, however, we found that after simulated blockade of RZ, but not TTX, *I*_*NaP*_ can be reactivated by transient hyperpolarization, due to differences between these drugs’ specific pharmacological mechanisms of action (Fig 6). This effect becomes critical in the intact respiratory network where these neurons receive strong, transient hyperpolarizing inhibition from post-I neurons during the expiratory phase of respiration. Correspondingly, our simulations of *I*_*NaP*_ blockade in the intact respiratory network predict that experimental application of RZ, but not TTX, will fail to effectively block *I*_*NaP*_ and respiratory rhythmicity.

### Insights into the role of *I*_*NaP*_ in the intact respiratory network

This computational study provides novel insight into the role of *I*_*NaP*_ in respiratory rhythmogenesis and pattern formation. In the intact respiratory network, the role of *I*_*NaP*_ is not well understood. Current thinking is that in the intact network *I*_*NaP*_ is largely inactivated and hence not essential for ongoing rhythm generation and pattern formation [17, 17]. This idea is based on computational modeling and the experimental observation that RZ application in the intact network fails to stop the respiratory rhythm, suggesting that active recruitment of *I*_*NaP*_ may not be essential to rhythmogenesis under these conditions [13].

This interpretation does not consider the dynamic interaction between network inhibition, tonic excitation and the voltage-dependent inactivation dynamics of *I*_*NaP*_, however. Moreover, this interpretation assumes that RZ effectively blocks *I*_*NaP*_ and overlooks the experimental observation that RZ application significantly reduces the amplitude of the respiratory rhythm generated by the intact network [13]. Indeed, the latter observation is not consistent with the idea that *I*_*NaP*_ is largely inactivated, as blocking an inactivated current should have no effect on rhythm characteristics. Our simulations support an alternative view. Specifically, to generate comparable reductions in network amplitude from simulated RZ blockade based on a shift in the *I*_*NaP*_ inactivation curve and from simulated TTX blockade based on a reduction in *I*_*NaP*_ conductance, *I*_*NaP*_ must strongly contribute to pre-I population bursting. Importantly, this contribution only occurs if, during periods when pre-I neurons are not spiking, *I*_*NaP*_ strongly deinactivates and *h*_*NaP*_ levels are higher in the intact network than in the isolated pre-I population (Figs 12 & 13).

An important finding of this study is the conclusion that *I*_*NaP*_ plays a critical role in respiratory rhythmogenesis, which is indicated by the prediction that complete blockade of *I*_*NaP*_ will stop respiratory rhythm generation in the intact network. Importantly, this finding comes to light only by considering the distinct pharmacological mechanisms of TTX and RZ in the context of the interaction between pre-I population excitability, inhibitory network interactions, and the associated dynamics of *I*_*NaP*_ inactivation.

We showed that the dependence of pre-I bursting on *I*_*NaP*_ in the intact network is a function of the excitability of the pre-I population (Fig 12 A). When inputs or neuromodulation sufficiently lowers the excitability of pre-I neurons, *I*_*NaP*_ becomes a necessity for burst dynamics, whereas with heightened excitability, bursting is dependent on inhibitory network interactions. While these mechanisms can act separately, there is a range of excitabilities for which pre-I bursting depends on both *I*_*NaP*_ and inhibitory network interactions.

In the intact network, the pre-I population receives transient inhibition from post-I neurons during the expiratory phase of respiration [13, 17, 18, 19, 20], which, our results show, may compromise the ability of RZ to effectively block *I*_*NaP*_. Our RZ simulations, tuned to match experimental data (specifically, a large reduction in pre-I output amplitude and no effect on frequency resulting from RZ application), suggest that the excitability of the pre-I population in the intact network is in a regime where rhythm generation depends on both *I*_*NaP*_ and inhibitory network interactions (Fig 12 A & 13). In these simulations, since RZ blockade of *I*_*NaP*_ fails to stop the rhythm and pre-I bursting is *I*_*NaP*_ -dependent, this indicates that RZ fails to completely block *I*_*NaP*_. In contrast, our simulations predict that under these network conditions a complete blockade of *I*_*NaP*_ by TTX will stop the respiratory rhythm. Importantly, at low concentrations (*≤*20 *nM*), TTX has been shown to selectively block *I*_*NaP*_ without affecting the action potential generating fast Na^+^ current [9]. Therefore, this prediction can be experimentally tested via bilateral microinfusion of TTX at low concentration into the pre-BötC in the intact preparation. If confirmed, this finding would illustrate the critical role of *I*_*NaP*_ in rhythm generation within intact respiratory circuits.

One caveat here is that the extent to which rhythm generation in the intact network relies on *I*_*NaP*_ and on inhibitory network interactions is highly dependent on the excitability of the pre-I population (Fig 12). Our simulations, tuned to match the results of [13], suggest that the excitability in the pre-I population must be close to the border between a regime where pre-I bursting depends exclusively on *I*_*NaP*_ and a regime where pre-I bursting depends on both *I*_*NaP*_ and inhibitory network interactions (Fig 12, 13). Therefore, with reasonable levels of variability, complete block of synaptic inhibition under these network conditions will inconsistently stop pre-I population oscillations and in instances where oscillations are stopped, the pre-I population will transition to a tonic mode of activity. Indeed, the prediction of proximity to the border is consistent with recent experimental data [18], which showed that simultaneous block/attenuation of GABAergic and glycinergic synaptic inhibition within the pre-BötC failed to abolish rhythmic phrenic nerve output in 83% experiments, and in experiments where oscillations were stopped, phrenic nerve activity culminated in tonic activity. Conditions that increase the excitability state of the pre-I population may transition the intact network into a regime where rhythm generation is no longer *I*_*NaP*_ -dependent, and complete *I*_*NaP*_ blockade under these conditions is not predicted to stop rhythmic oscillations. Hypoxia and hypercapnia are perturbations that are likely to affect pre-I excitability and consequently the role of *I*_*NaP*_ in the intact network.

It is important to note that the model used in the current study is the same as that considered in a past work that came to some different conclusions [13]. Specifically, the authors of that work suggested that *I*_*NaP*_ is largely inactivated and hence is not critical for rhythm generation in the intact network. This previous study, however, did not systematically consider the interaction between pre-I population excitability, inhibitory network interactions, and the resulting dynamics of *I*_*NaP*_ inhibition. It also did not consider the distinct pharmacological mechanism of action of RZ and block was simulated by reducing *g*_*NaP*_. With the parameter set used for simulations representing the intact network in [13], the pre-I population excitability was set at *g*_*tonic*_ = 0.98 *ns* and the strength of post-I to pre-I inhibition was 0.225 *nS*. In [13] and our simulations, *I*_*NaP*_ is largely inactivated under these conditions. Consequently, under these parameters, simulated TTX or RZ block of *I*_*NaP*_ cannot capture the large reduction in amplitude seen with experimental RZ application in the intact network. Therefore, conclusions drawn from these simulations about the excitability state of the pre-I population, *I*_*NaP*_ inactivation dynamics, and the role of *I*_*NaP*_ in rhythm generation in the intact respiratory network may not accurately represent the underlying mechanisms, dynamics, and conditions in these experimental preparations.

### RZ-dependent reduction in excitatory synaptic transmission

Characterizing the off-target effects of RZ in respiratory circuits may be critical for understanding how RZ application impacts pre-I network dynamics [25]. Comparison of experimental and simulated data in this study supports the idea that RZ (but not TTX) application not only alters *I*_*NaP*_ but also attenuates excitatory synaptic transmission. In regions outside of the pre-BötC, RZ has been shown to block excitatory synaptic transmission at doses comparable to those used in respiratory circuits (0*-*20 *µm*) [26, 26]. The effects of RZ on synaptic transmission or other off-target effects have not been characterized in the pre-BötC, however. Experimentally, bilateral microinfusion of TTX or RZ into the pre-BötC stops the fictive respiratory rhythm in in vitro slice preparations. Analysis of the time course of these drugs reveals that before rhythm termination, RZ reduces the frequency and amplitude of integrated hypoglossal nerve motor output, whereas TTX only reduces frequency [9].

In our simulations of the isolated pre-I network, selective blockade of *I*_*NaP*_ by either the pharmacological mechanism of TTX or RZ reduces network frequency without affecting amplitude (Fig 8A, B). In contrast, blockade of excitatory synaptic transmission selectively reduces network amplitude with little effect on frequency (Fig 8C), which is consistent with experimental data [23]. Therefore, to account for the reduction in amplitude seen with experimental RZ application, our results suggest that RZ must reduce the strength of the excitatory synaptic transmission in addition to modulating *I*_*NaP*_ (Fig 9). The prediction that RZ attenuates *I*_*SynE*_ is testable by isolating the synaptic current in vitro using voltage-clamp recordings in conjunction with RZ and TTX application.

Importantly, although RZ blocks both *I*_*NaP*_ and *I*_*SynE*_ in our simulations of the isolated pre-I network, rhythm generation in this system is abolished due to the reduction of *I*_*NaP*_, not *I*_*SynE*_. Partial reduction of *I*_*SynE*_ alone is not sufficient to stop rhythm generation in our simulations (Fig 8C). These results support the hypothesis that *I*_*NaP*_ is critical for rhythm generation in the pre-BötC in in vitro slice preparations.

It is important to mention that additional off-target effects of RZ have been reported in neurons outside of respiratory circuits such as: potentiation of calcium-dependent K^+^ current, inhibition of fast Na^+^ current, inhibition of voltage-gated Ca^2+^ current, inhibition of voltage-gated K^+^ current, and inhibition of the glutimate receptor N-methyl-D-aspartate receptor (NMDA) [25, 25]. Detailed consideration of these additional off-target RZ effects are are beyond the scope of this study. Many of these off-target effects occur at relatively high RZ concentrations and are therefore not likely to effect the results of this study.

Note that our simulations of *I*_*NaP*_ block in the isolated pre-I population are specifically tuned to match data where TTX or RZ application is delivered via bilateral microinfusion into the *I*_*NaP*_, as opposed to bath application [9]. This choice was made because bilateral microinfusion is likely a more targeted/localized approach and avoids the issues of non-uniform and off-target drug effects that may arise with bath application.

### Non-uniform pharmacological blockade in the pre-BötC pre-I population

The role of *I*_*NaP*_ in respiratory rhythm generation in the isolated pre-I population is also unclear and a highly debated topic within the field. This debate is fueled by the diverse and seemingly contradictory effects of *I*_*NaP*_ block by bath application of TTX or RZ in in vitro slice preparations. For example, [9] found that in vitro bath application of either TTX or RZ reliably decreased the frequency and to a lesser extent the amplitude of network bursting before oscillations abruptly stopped. In contrast, [32, 32] found that bath application of RZ at comparable concentrations resulted in a large reduction in the network amplitude but had no effect on or even increased frequency and failed to stop network oscillations. Consideration of non-uniform *I*_*NaP*_ block may provide an explanation for these seemingly contradictory findings.

With bath application of TTX or RZ, block of *I*_*NaP*_ is unlikely to be uniform since drug penetration is dependent on passive diffusion. Therefore, with bath application, *I*_*NaP*_ block will affect neurons close to the surface of the slice first and progress towards neurons in the center. In Fig 10 we show that the result of progressive *I*_*NaP*_ block across the isolated pre-I population is highly dependent on the dynamics of the neurons (pacemaker vs non-pacemaker) affected. Our simulations can explain the diversity of experimental results if we consider the effects of slice thickness and assume that pacemaker neurons are preferentially located near the center of the larger population of pre-I neurons; see Fig 14.

**Fig 14.**
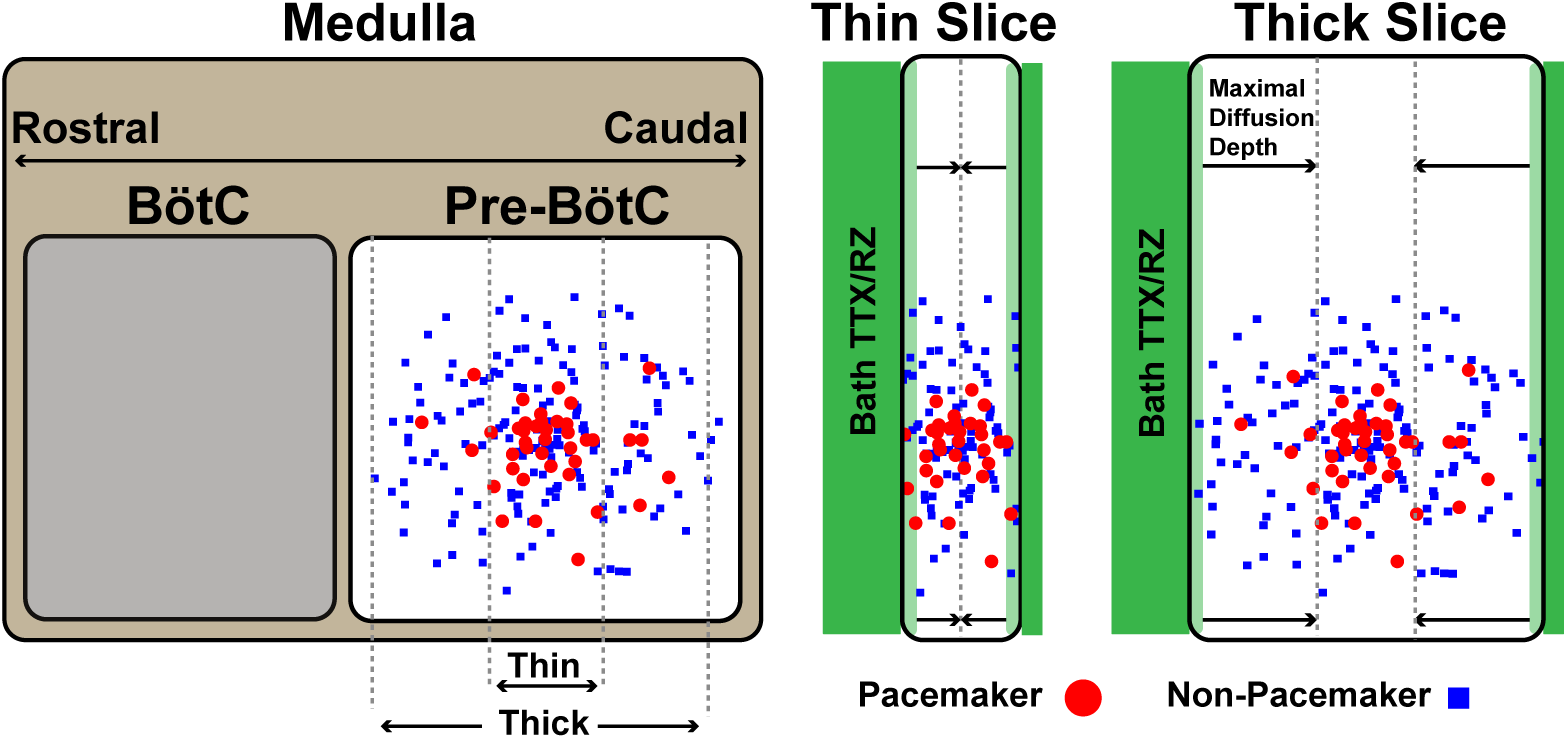
Sketch of proposed distribution of pacemaker and non-pacemaker neurons within the pre-BötC as well as the the direction and depth of TTX and RZ penetration in thick and thin in vitro slice preparation containing the pre-BötC. The direction of diffusion and maximal diffusion depth in the thin and thick slice sketches are indicated by arrows and vertical dashed lines. Notice that in a thick slice non-pacemaker neurons are effected first and TTX and RZ fail to fully penetrate the thick slice and in this sketch.

In a thin slice, both pacemaker and non-pacemaker neurons are located close to the surface and an applied drug should fully diffuse through the slice. In this case, as TTX or RZ diffuses in, *I*_*NaP*_ will be blocked in both pacemaker and non-pacemaker neurons, which is predicted to affect both amplitude and frequency until oscillations eventually stop (Fig 10C). In contrast, with a thick slice, TTX and RZ may not diffuse all the way to the center and the neurons affected first would primarily be the peripheral non-pacemaker neurons. With *I*_*NaP*_ blocked predominantly in non-pacemaker neurons, our results predict an effect on the amplitude of oscillations (Fig 10B). If the drug diffuses deeper into the slice, both pacemaker and non-pacemaker neurons will be impacted, which will further decrease amplitude and start to decrease frequency (Fig 10C).

Therefore, our model predicts that in the case of a thick slice, the lack of frequency effects and the failure to stop network oscillations derive from incomplete/inadequate drug penetration into the slice. It is unknown how far TTX or RZ diffuse into neuronal tissue. However, a straightforward prediction of incomplete drug penetration is the observation of pacemaker neurons that appear insensitive to *I*_*NaP*_ block. Consistent with this prediction, [24] found pacemaker neurons that are insensitive both to RZ application and to Cd^2+^ application that blocks Ca^2+^ dynamics thought to underlie rhythmic properties in some pacemaker neurons. An experimental preparation that contains the same population of neurons and likely avoids the issue of non-uniform/incomplete *I*_*NaP*_ block is an in situ rat brain stem-spinal cord preparation where where regions rostal to the pre-BötC are removed and RZ is delivered by arterial perfusion. In this preparation, application of RZ at concentrations greater than 10 *µm* consistently stopped rhythmic oscillations of the pre-I population [13].

### Limitations of this study

In the pre-BötC, coupling of a calcium-activated non-selective cation current (*I*_*CAN*_) and intracellular calcium transients is proposed to represent a biophysical mechanism underlying rhythmogenesis at the cellular- and/or network-level [24, 33, 34, 35, 36, 37, 38, 39]. For simplicity, however, explicit representation of *I*_*CAN*_ and intracellular calcium dynamics were omitted in this study. This choice can be justified by considering the effects of pharmacological block of *I*_*CAN*_ in conjunction with recent data-driven computational work. In in vitro slice preparations containing the pre-BötC, pharmacological blockade of *I*_*CAN*_ results in a large reduction in the amplitude of network oscillation with no or minor perturbation of frequency, suggesting that the primary role of *I*_*CAN*_ in these circuits is amplitude generation/regulation [40, 40]. A recent data-driven computational study that included *I*_*NaP*_, *I*_*CAN*_, and intracellular calcium dynamics [41] found that in order to match experimental data, *I*_*CAN*_ activation must be strongly coupled to synaptically triggered calcium transients, as suggested by [35, 37, 38, 39, 42]. Consequently, *I*_*CAN*_ acts as a mechanism that amplifies excitatory synaptic inputs within the pre-BötC. As such, *I*_*CAN*_ can be treated as an excitatory post synaptic current that, in our model, could correspond to a portion of the total excitatory current *I*_*SynE*_. Consistent with this idea, blockade of *I*_*SynE*_ in our model of the isolated pre-BötC is consistent with the effects seen with experimental blockade of *I*_*CAN*_ (Fig 8C).

Another issue that we do not explore in this work is the contribution of additional burst termination mechanisms. At the neuronal and population level, mechanisms of burst termination are critical for the generation of ongoing rhythmic oscillations. In the current model, burst termination is exclusively dependent on the inactivation of *I*_*NaP*_. However, in respiratory neurons and circuits, additional mechanisms of burst termination have been proposed, such as slowly activating potassium channels [14], inositol triphosphate (IP3) effects [36, 43, 44], Na^+^/K^+^ ATPase electrogenic pumps [45, 45], and synaptic depression [46, 46], for example. Inclusion of additional burst terminating mechanisms in *I*_*NaP*_ -dependent rhythmogenic models of pre-BötC neurons increases the dynamic range where bursting occurs, making rhythm generation more robust [44], and may change the effects of *I*_*NaP*_ block on neuronal activity in these models. For example, inclusion of a Na^+^/K^+^ ATPase electrogenic pump in [44], in addition to *I*_*NaP*_, dramatically increases the dynamic range where *I*_*NaP*_ block transitions the neuron from a bursting to a tonic mode of activity, as opposed to a transition to a silent model of activity. Although theoretically interesting, additional burst termination mechanisms were not necessary to match a wide range of experimental data in our simulations and therefore were not included in this study.

Several additional properties may modulate pre-BötC oscillations but are not considered in this work. Local inhibition within the pre-BötC, for example, may play a role in pre-I burst synchrony, variability and patterning [20], but is not critical for intrinsic pre-BötC oscillations [47]. Similarly, spatial variability in network organization and synaptic connection probability as well as stochasticity and neuromodulation may play important roles in shaping pattern formation in respiratory circuits [20, 37, 48, 49, 50, 51], but consideration of these additional features is beyond the scope of this study and is left for future work.

### Broader implications

*I*_*NaP*_ is not unique to respiratory circuits. Experimental and computational results have suggested an active role for *I*_*NaP*_ in shaping activity patterns of putative locomotor CPG neurons [52, 53, 54] and in contributing to locomotor rhythm generation [53, 55, 56]. Similar roles are supported for *I*_*NaP*_ in generating the neural rhythm underlying mastication [58, 58]. *I*_*NaP*_ is also expressed and contributes to neural activity elsewhere in the central nervous system [60, 60]. RZ and TTX have been used in experimental investigations of these neurons and their activity patterns. Additionally, RZ has been considered as a neuroprotective or anticonvulsant agent [8] and is actively investigated for its for therapeutic potential in the treatment of multiple diseases such as obsessive–compulsive disorder (OCD) [61, 62, 63], anxiety [65, 65], depression [67, 67], multiple sclerosis (MS) [68], and amyotrophic lateral sclerosis (ALS) [69]. Therefore, understanding RZ’s pharmacological mechanisms of action may be important for understanding its beneficial effects in these disease states.

## Conclusion

This computational study, in conjunction with experimental data, has illustrated the general importance of considering the pharmacological mechanism(s) of action when interpreting and simulating data from pharmacological blockade studies. In the case of respiratory circuits, we show that (1) RZ may fail to effectively block *I*_*NaP*_ in the intact network due the specific pharmacological mechanism of action and transient inhibitory network interactions, (2) pre-I bursting in the intact network is likely dependent on both inhibitory network interactions and *I*_*NaP*_, and (3) experimental TTX application in the intact network is predicted to terminate respiratory rhythmicity. These findings suggest that *I*_*NaP*_ is less inactivated than has been previously proposed and that it plays a critical role in respiratory rhythm generation under in vivo conditions. If experimentally confirmed, these predictions will advance our understanding of the mechanisms of rhythm generation in brainstem respiratory circuits. These findings also have implications in understanding the role of *I*_*NaP*_ in CPGs other than respiratory circuits, in elucidating the effects of RZ across the CNS in the treatment of various conditions, and in interpreting and simulating data from pharmacological blockade studies in general.

## Materials and methods

### Model description

Neurons were simulated with single compartment models incorporating Hodgkin-Huxley style conductances based on previously described models [13, 14, 44]. The membrane potential *V*_*m*_ for each neuron is given by the following differential equation:

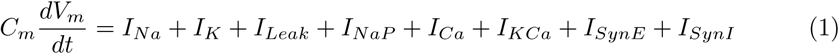

where *C*_*m*_ = 36.0 *pF* is the cell capacitance, *I*_*Na*_, *I*_*K*_, *I*_*Leak*_, *I*_*NaP*_, *I*_*Ca*_, *I*_*KCa*_, *I*_*SynE*_, and *I*_*SynI*_ are the sodium, potassium, leak, persistent sodium, high-voltage activated calcium, calcium-activated potassium, excitatory synaptic and inhibitory synaptic ionic currents, respectively. The currents are defined as follows:

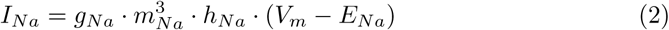

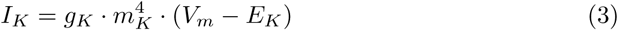

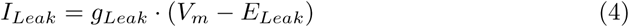

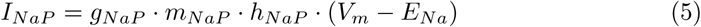

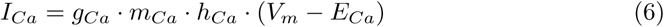

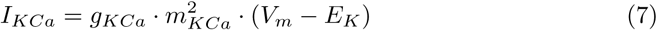

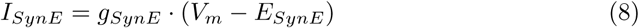

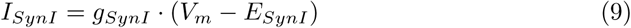

where *g*_*i*_ is the maximum conductance, *E*_*i*_ is the reversal potential, and *m*_*i*_ and *h*_*i*_ are gating variables for channel activation and inactivation for current *I*_*i*_. Values used for *g*_*i*_ and *E*_*i*_ are given in Table 1 and 2.

**Table 1.**
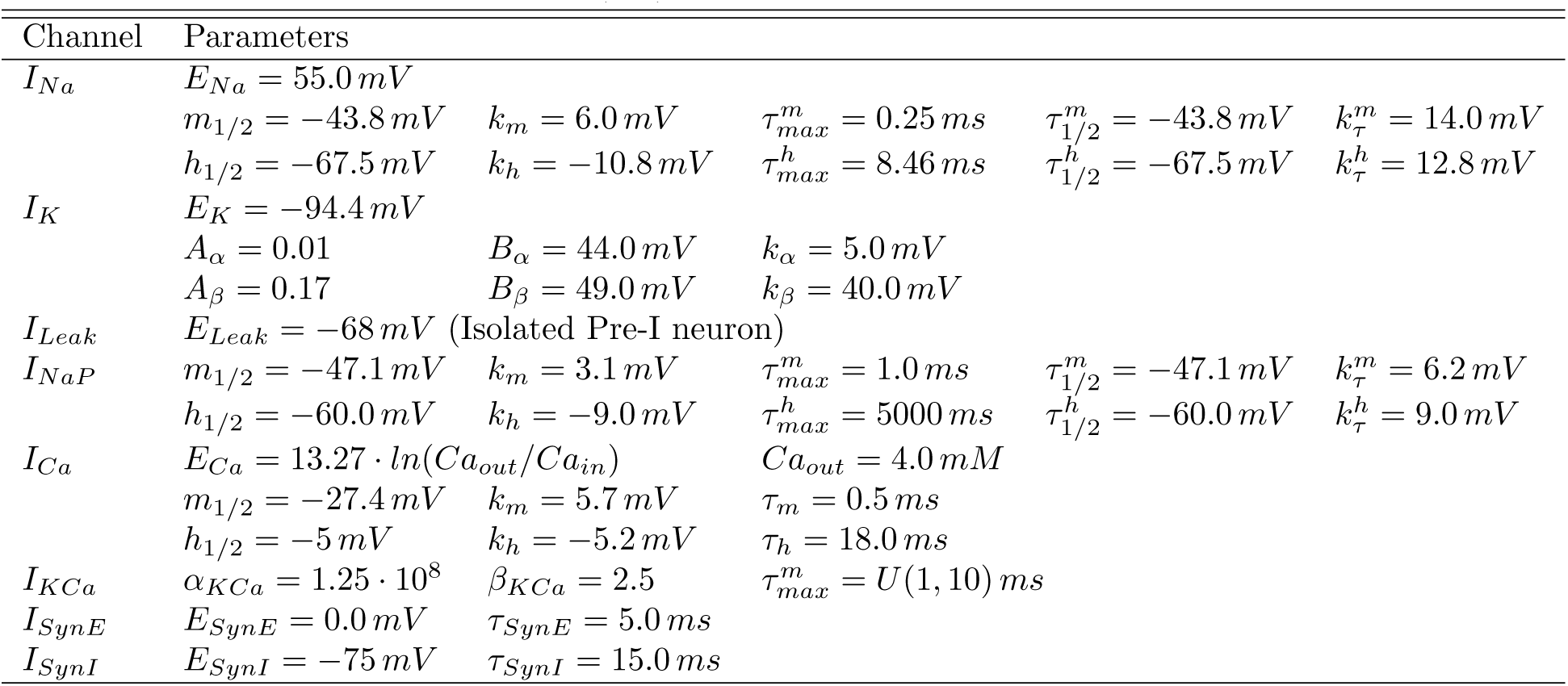
Ionic Channel Parameters. *U* (*a, b*) indicates a uniform distribution from *a* to *b*.

**Table 2.** Neuronal Type Specific Parameters. *U* (*a, b*) indicates a uniform distribution from *a* to *b*.

| Type | $g_{Na} (nS)$ | $g_K (nS)$ | $g_{Leak} (nS)$ | $E_{Leak} (mV)$ | $g_{NaP} (nS)$ | $g_{Ca} (nS)$ | $g_{KCa} (nS)$ | $g_{TonicE} (nS)$ |
| --- | --- | --- | --- | --- | --- | --- | --- | --- |
| Pre-I | 170 | 180 | 2.25 | $U(-66.5, -69.5)$ | 5.0 | - | - | $0 - 0.5$ |
| Early-I | 400 | 250 | 6.0 | $U(-58.8, -61.2)$ | - | 0.34 | $U(3.0, 6.0)$ | 0.25 |
| Aug-E | 400 | 250 | 6.0 | $U(-58.8, -61.2)$ | - | 0.042 | $U(3.0, 6.0)$ | 0.475 |
| Post-I | 400 | 250 | 6.0 | $U(-58.8, -61.2)$ | - | 0.17 | $U(3.0, 6.0)$ | 0.3 |

**Table 3.** Maximal weight of excitatory (*W*_*MaxE*_) and inhibitory (*W*_*MaxI*_) synaptic connections in nS.

| | | $W_{MaxE}$ Target (i) | | | | $W_{MaxI}$ Target (i) | | | |
| --- | --- | --- | --- | --- | --- | --- | --- | --- | --- |
|  |  | Pre-I | Early-I | Aug-E | Post-I | Pre-I | Early-I | Aug-E | Post-I |
| source (j) | Pre-I | 0.03 | 0.125 | - | - | - | - | - | - |
|  | Early-I | - | - | - | - | - | - | 0.22 | 0.02635 |
|  | Aug-E | - | - | - | - | 0.0075 | 0.05 | - | 0.01125 |
|  | Post-I | - | - | - | - | 0.75 | 0.372 | 0.15 | - |

Dynamics of the gating variables *m, h* for all channels are described by the following differential equation:

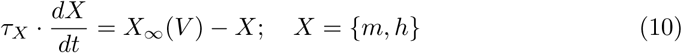

where *X*_*∞*_ represents steady-state activation/inactivation and *τ*_*X*_ is a time constant. For *I*_*Na*_, *I*_*NaP*_, and *I*_*Ca*_, the functions *X*_*∞*_ and *τ*_*X*_ take the forms

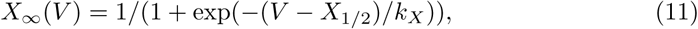

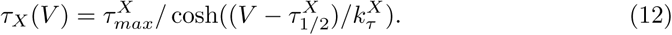

The values of the parameters 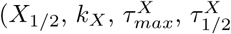, and 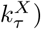) corresponding to *I*_*Na*_ *I*_*NaP*_ and *I*_*Ca*_ are given in Table 1.

For the voltage-gated potassium channel, steady-state activation 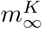 and time constant 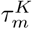 given by the expressions

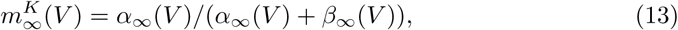

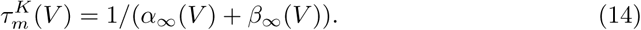

Where

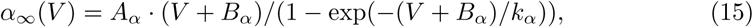

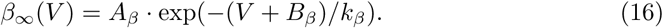

The values for the constants *A*_*α*_, *A*_*β*_, *B*_*α*_, *B*_*β*_, *k*_*α*_, and *k*_*β*_ are also given in Table 1.

For *I*_*KCa*_, the steady-state activation 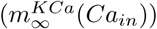 and time constant 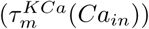 depend on the intracellular *Ca*^2+^ concentration (*Ca*_*in*_) as follows:

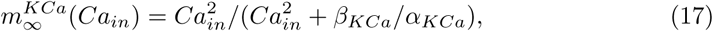

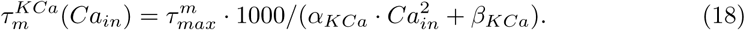

The values of *α*_*KCa*_, *β*_*KCa*_ and 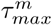 can be found in Table 1.

*Ca*_*in*_ is determined by the balance of *Ca*^2+^ influx carried by *I*_*Ca*_ and efflux via *Ca*^2+^ pump. The dynamics of *Ca*_*in*_ is described as follows:

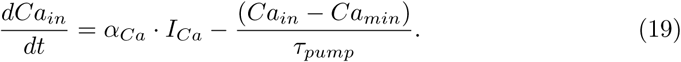

where *α*_*Ca*_ = 2.5 10^*-*5^ *mM/f C* is a conversion factor relating current to rate of change of *Ca*_*in*_, *τ*_*pump*_ = 500 *ms* is the time constant for the *Ca*^2+^ pump and *Ca*_*min*_ = 5.0. 10^*-*6^ *mM* is a minimal baseline calcium concentration.

The total synaptic conductances 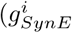 and 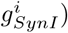 of the *i*^*th*^ target neuron are described by the following equation:

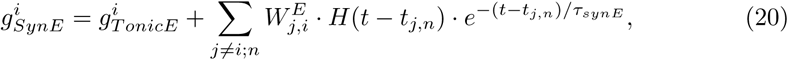

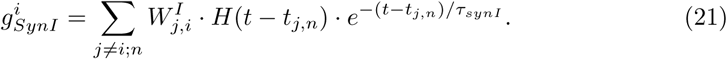

Where 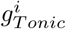 is the tonic excitatory conductance, 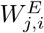 and 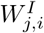 are the weights of the excitatory and inhibitory synaptic connection from the source neuron *j* to the target neuron *i, H*(.) is the Heaviside step function, and *t* denotes time. *τ*_*SynE*_ and *τ*_*SynI*_ are exponential decay constants for excitatory and inhibitory synapses. *t*_*j,n*_ is the time at which the *n*^*th*^ action potential is generated in neuron *j* and reaches neuron *i*. The weights of excitatory and inhibitory conductances were uniformly distributed such that 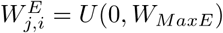 and 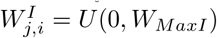 where *W*_*MaxE*_ and *W*_*MaxI*_ are constants. Parameter values are listed in Tables 1 & 3.

### Network construction

The simulations representing the intact respiratory network and the isolated pre-I population were reconstructed from the model description used in [13]. In the intact network each population (pre-I, early-I, aug-E, post-I) consists of 50 model neurons governed by the equations presented in the previous subsection, with intrinsic neuronal parameter values given in Table 2. Note that heterogeneity was introduced by uniformly distributing the parameters *E*_*leak*_ within each population as well as the weights of excitatory 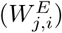 and inhibitory 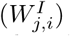 synaptic connections; see Tables 2 & 3. The tonic excitatory drive (*g*_*Tonic*_) within the isolated pre-I population model (meant to represent in vitro slice preparations) was tuned such that the percentage of bursting neurons in the synaptically decoupled network is between 20-30%, consistent with experimental observations [24]. The maximal weight (*W*_*MaxE*_) of excitatory synapses between neurons was then tuned to achieve complete network synchronization, where all neurons are active during network oscillations. In full network simulations (meant to represent in vivo or in situ brain stem-spinal cord preparations), *g*_*Tonic*_ was treated as a variable to match experimental observations [13].

### Data analysis and definitions

Data generated from simulations was post-processed in Matlab (Mathworks, Inc.). An action potential was defined to have occurred in a neuron when its membrane potential *V*_*m*_ increased through –35*mV*. Histograms of population activity were calculated as the number of action potentials per 50 *ms* bin per neuron with units of *APs/*(*s · neuron*). Population amplitude and frequency were calculated by identifying the peaks and the inverse of the interpeak interval from the population histograms.

### Integration methods

All simulations were performed locally on an 8-core Linux-based operating system. Simulation software was custom written in C++. Numerical integration was performed using the first-order Euler method with a fixed step-size (Δ*t*) of 0.025*ms*.

## Acknowledgments

We thank Dr. Jeffrey Smith for providing experimental data. This study was partially supported by the NSF via award DMS 1612913 and the CRCNS award DMS 1724240.

## References

1. Cestéle S, Catterall WA. Molecular mechanisms of neurotoxin action on voltage-gated sodium channels. Biochimie. 2000;82(9-10):883–892.

2. Stevens M, Peigneur S, Tytgat J. Neurotoxins and their binding areas on voltage-gated sodium channels. Frontiers in pharmacology. 2011;2:71.

3. Hille B. Ionic Channels of Excitable Membranes. 1992;.

4. Hebert T, Drapeau P, Pradier L, Dunn RJ. Block of the rat brain IIA sodium channel alpha subunit by the neuroprotective drug riluzole. Molecular pharmacology. 1994;45(5):1055–1060.

5. Song JH, Huang CS, Nagata K, Yeh JZ, Narahashi T. Differential action of riluzole on tetrodotoxin-sensitive and tetrodotoxin-resistant sodium channels. Journal of Pharmacology and Experimental Therapeutics. 1997;282(2):707–714.

6. Ptak K, Zummo GG, Alheid GF, Tkatch T, Surmeier DJ, McCrimmon DR. Sodium currents in medullary neurons isolated from the pre-Bötzinger complex region. Journal of Neuroscience. 2005;25(21):5159–5170.

7. Benoit E, Escande D. Riluzole specifically blocks inactivated Na channels in myelinated nerve fibre. Pflügers Archiv. 1991;419(6):603–609.

8. Urbani A, Belluzzi O. Riluzole inhibits the persistent sodium current in mammalian CNS neurons. European Journal of Neuroscience. 2000;12(10):3567–3574.

9. Koizumi H, Smith JC. Persistent Na+ and K+-dominated leak currents contribute to respiratory rhythm generation in the pre-Bötzinger complex in vitro. Journal of Neuroscience. 2008;28(7):1773–1785.

10. Bellingham MC. A review of the neural mechanisms of action and clinical efficiency of riluzole in treating amyotrophic lateral sclerosis: what have we learned in the last decade? CNS neuroscience & therapeutics. 2011;17(1):4–31.

11. MacIver MB, Amagasu SM, Mikulec AA, Monroe FA. Riluzole AnesthesiaUse-Dependent Block of Presynaptic Glutamate Fibers. Anesthesiology: The Journal of the American Society of Anesthesiologists. 1996;85(3):626–634.

12. Del Negro CA, Koshiya N, Butera Jr RJ, Smith JC. Persistent sodium current, membrane properties and bursting behavior of pre-botzinger complex inspiratory neurons in vitro. Journal of neurophysiology. 2002;88(5):2242–2250.

13. Smith JC, Abdala A, Koizumi H, Rybak IA, Paton JF. Spatial and functional architecture of the mammalian brain stem respiratory network: a hierarchy of three oscillatory mechanisms. Journal of neurophysiology. 2007;98(6):3370–3387.

14. Butera Jr RJ, Rinzel J, Smith JC. Models of respiratory rhythm generation in the pre-Botzinger complex. I. Bursting pacemaker neurons. Journal of neurophysiology. 1999;82(1):382–397.

15. Butera Jr RJ, Rinzel J, Smith JC. Models of respiratory rhythm generation in the pre-Botzinger complex. II. Populations of coupled pacemaker neurons. Journal of neurophysiology. 1999;82(1):398–415.

16. Smith J. Integration of cellular and network mechanisms in mammalian oscillatory motor circuits: insights from the respiratory oscillator. Neurons, Networks, and Motor Behavior. 1997; p. 97–104.

17. Richter DW, Smith JC. Respiratory rhythm generation in vivo. Physiology. 2014;29(1):58–71.

18. Marchenko V, Koizumi H, Mosher B, Koshiya N, Tariq MF, Bezdudnaya TG, et al. Perturbations of respiratory rhythm and pattern by disrupting synaptic inhibition within pre-Bötzinger and Bötzinger complexes. eNeuro. 2016; p. ENEURO–0011.

19. Cregg JM, Chu KA, Dick TE, Landmesser LT, Silver J. Phasic inhibition as a mechanism for generation of rapid respiratory rhythms. Proceedings of the National Academy of Sciences. 2017;114(48):12815–12820.

20. Baertsch NA, Baertsch HC, Ramirez JM. The interdependence of excitation and inhibition for the control of dynamic breathing rhythms. Nature communications. 2018;9(1):843.

21. Bertram R, Rubin JE. Multi-timescale systems and fast-slow analysis. Mathematical biosciences. 2017;287:105–121.

22. Smith JC, Ellenberger HH, Ballanyi K, Richter DW, Feldman JL. Pre-Botzinger complex: a brainstem region that may generate respiratory rhythm in mammals. Science. 1991;254(5032):726–729.

23. Johnson SM, Smith JC, Funk GD, Feldman JL. Pacemaker behavior of respiratory neurons in medullary slices from neonatal rat. Journal of neurophysiology. 1994;72(6):2598–2608.

24. Peña F, Parkis MA, Tryba AK, Ramirez JM. Differential contribution of pacemaker properties to the generation of respiratory rhythms during normoxia and hypoxia. Neuron. 2004;43(1):105–117.

25. Morgado-Valle C, Beltran-Parrazal L. Respiratory rhythm generation: the whole is greater than the sum of the parts. In: The Plastic Brain. Springer; 2017. p. 147–161.

26. Bellingham MC. Pre-and postsynaptic mechanisms underlying inhibition of hypoglossal motor neuron excitability by riluzole. Journal of neurophysiology. 2013;110(5):1047–1061.

27. Rybak IA, Abdala AP, Markin SN, Paton JF, Smith JC. Spatial organization and state-dependent mechanisms for respiratory rhythm and pattern generation. Progress in brain research. 2007;165:201–220.

28. Rubin JE, Shevtsova NA, Ermentrout GB, Smith JC, Rybak IA. Multiple rhythmic states in a model of the respiratory central pattern generator. Journal of Neurophysiology. 2009;101(4):2146–2165.

29. Molkov YI, Rubin JE, Rybak IA, Smith JC. Computational models of the neural control of breathing. Wiley Interdisciplinary Reviews: Systems Biology and Medicine. 2017;9(2):e1371.

30. Fortuna MG, Kügler S, Hülsmann S. Probing the function of glycinergic neurons in the mouse respiratory network using optogenetics. Respiratory physiology & neurobiology. 2018;.

31. Diekman CO, Thomas PJ, Wilson CG. Eupnea, tachypnea, and autoresuscitation in a closed-loop respiratory control model. Journal of Neurophysiology. 2017;118(4):2194–2215.

32. Del Negro CA, Morgado-Valle C, Feldman JL. Respiratory rhythm: an emergent network property? Neuron. 2002;34(5):821–830.

33. Thoby-Brisson M, Ramirez JM. Identification of two types of inspiratory pacemaker neurons in the isolated respiratory neural network of mice. Journal of neurophysiology. 2001;86(1):104–112.

34. Del Negro CA, Morgado-Valle C, Hayes JA, Mackay DD, Pace RW, Crowder EA, et al. Sodium and calcium current-mediated pacemaker neurons and respiratory rhythm generation. Journal of Neuroscience. 2005;25(2):446–453.

35. Feldman JL, Del Negro CA. Looking for inspiration: new perspectives on respiratory rhythm. Nature Reviews Neuroscience. 2006;7(3):232.

36. Pace RW, Mackay DD, Feldman JL, Del Negro CA. Inspiratory bursts in the preBötzinger complex depend on a calcium-activated non-specific cation current linked to glutamate receptors in neonatal mice. The Journal of physiology. 2007;582(1):113–125.

37. Mironov S. Metabotropic glutamate receptors activate dendritic calcium waves and TRPM channels which drive rhythmic respiratory patterns in mice. The Journal of physiology. 2008;586(9):2277–2291.

38. Rubin JE, Hayes JA, Mendenhall JL, Del Negro CA. Calcium-activated nonspecific cation current and synaptic depression promote network-dependent burst oscillations. Proceedings of the National Academy of Sciences. 2009;106(8):2939–2944.

39. Del Negro CA, Hayes JA, Pace RW, Brush BR, Teruyama R, Feldman JL. Synaptically activated burst-generating conductances may underlie a group-pacemaker mechanism for respiratory rhythm generation in mammals. In: Progress in brain research. vol. 187. Elsevier; 2010. p. 111–136.

40. Koizumi H, John TT, Chia JX, Tariq MF, Phillips RS, Mosher B, et al. Transient Receptor Potential Channels TRPM4 and TRPC3 Critically Contribute to Respiratory Motor Pattern Formation but not Rhythmogenesis in Rodent Brainstem Circuits. eNeuro. 2018; p. ENEURO–0332.

41. Phillips R, John TT, Koizumi H, Molkov YI, Smith JC. Biophysical mechanisms in the mammalian respiratory oscillator re-examined with a new data-driven computational model. bioRxiv. 2018; p. 415190.

42. Morgado-Valle C, Beltran-Parrazal L, DiFranco M, Vergara JL, Feldman JL. Somatic Ca2+ transients do not contribute to inspiratory drive in preBötzinger complex neurons. The Journal of physiology. 2008;586(18):4531–4540.

43. Pace RW, Del Negro CA. AMPA and metabotropic glutamate receptors cooperatively generate inspiratory-like depolarization in mouse respiratory neurons in vitro. European Journal of Neuroscience. 2008;28(12):2434–2442.

44. Jasinski PE, Molkov YI, Shevtsova NA, Smith JC, Rybak IA. Sodium and calcium mechanisms of rhythmic bursting in excitatory neural networks of the pre-B ötzinger complex: a computational modelling study. European Journal of Neuroscience. 2013;37(2):212–230.

45. Krey RA, Goodreau AM, Arnold TB, Del Negro CA. Outward currents contributing to inspiratory burst termination in preBötzinger complex neurons of neonatal mice studied in vitro. Frontiers in neural circuits. 2010;4:124.

46. Kottick A, Del Negro CA. Synaptic depression influences inspiratory–expiratory phase transition in Dbx1 interneurons of the preBötzinger complex in neonatal mice. Journal of Neuroscience. 2015;35(33):11606–11611.

47. Shao XM, Feldman JL. Respiratory rhythm generation and synaptic inhibition of expiratory neurons in pre-Botzinger complex: differential roles of glycinergic and GABAergic neural transmission. Journal of neurophysiology. 1997;77(4):1853–1860.

48. Nesse WH, Del Negro CA, Bressloff PC. Oscillation regularity in noise-driven excitable systems with multi-time-scale adaptation. Physical review letters. 2008;101(8):088101.

49. Carroll MS, Ramirez JM. Cycle-by-cycle assembly of respiratory network activity is dynamic and stochastic. Journal of neurophysiology. 2012;109(2):296–305.

50. Carroll MS, Viemari JC, Ramirez JM. Patterns of inspiratory phase-dependent activity in the in vitro respiratory network. Journal of neurophysiology. 2012;109(2):285–295.

51. Yu H, Dhingra RR, Dick TE, Galan RF. Effects of ion-channel noise on neural circuits: an application to the respiratory pattern generator to investigate breathing variability. American Journal of Physiology-Heart and Circulatory Physiology. 2016;.

52. Tazerart S, Viemari JC, Darbon P, Vinay L, Brocard F. Contribution of persistent sodium current to locomotor pattern generation in neonatal rats. Journal of neurophysiology. 2007;98(2):613–628.

53. Tazerart S, Vinay L, Brocard F. The persistent sodium current generates pacemaker activities in the central pattern generator for locomotion and regulates the locomotor rhythm. Journal of Neuroscience. 2008;28(34):8577–8589.

54. Ziskind-Conhaim L, Wu L, Wiesner EP. Persistent sodium current contributes to induced voltage oscillations in locomotor-related hb9 interneurons in the mouse spinal cord. Journal of neurophysiology. 2008;100(4):2254–2264.

55. Brocard F, Tazerart S, Vinay L. Do pacemakers drive the central pattern generator for locomotion in mammals? The Neuroscientist. 2010;16(2):139–155.

56. Brocard F, Shevtsova NA, Bouhadfane M, Tazerart S, Heinemann U, Rybak IA, et al. Activity-dependent changes in extracellular Ca2+ and K+ reveal pacemakers in the spinal locomotor-related network. Neuron. 2013;77(6):1047–1054.

57. Brocard F, Verdier D, Arsenault I, Lund JP, Kolta A. Emergence of intrinsic bursting in trigeminal sensory neurons parallels the acquisition of mastication in weanling rats. Journal of neurophysiology. 2006;96(5):2410–2424.

58. Tsuruyama K, Hsiao CF, Chandler SH. Participation of a persistent sodium current and calcium-activated nonspecific cationic current to burst generation in trigeminal principal sensory neurons. Journal of neurophysiology. 2013;110(8):1903–1914.

59. Crill WE. Persistent sodium current in mammalian central neurons. Annual review of physiology. 1996;58(1):349–362.

60. van Drongelen W, Koch H, Elsen FP, Lee HC, Mrejeru A, Doren E, et al. Role of persistent sodium current in bursting activity of mouse neocortical networks in vitro. Journal of Neurophysiology. 2006;96(5):2564–2577.

61. Grant P, Song JY, Swedo SE. Review of the use of the glutamate antagonist riluzole in psychiatric disorders and a description of recent use in childhood obsessive-compulsive disorder. Journal of child and adolescent psychopharmacology. 2010;20(4):309–315.

62. Pittenger C, Kelmendi B, Wasylink S, Bloch MH, Coric V. Riluzole augmentation in treatment-refractory obsessive-compulsive disorder: a series of 13 cases, with long-term follow-up. Journal of clinical psychopharmacology. 2008;28(3):363–367.

63. Pittenger C, Bloch MH, Wasylink S, Billingslea E, Simpson R, Jakubovski E, et al. Riluzole augmentation in treatment-refractory obsessive-compulsive disorder: a pilot placebo-controlled trial. The Journal of clinical psychiatry. 2015;76(8):1075.

64. Mathew SJ, Amiel JM, Coplan JD, Fitterling HA, Sackeim HA, Gorman JM. Open-label trial of riluzole in generalized anxiety disorder. American Journal of Psychiatry. 2005;162(12):2379–2381.

65. Pittenger C, Coric V, Banasr M, Bloch M, Krystal JH, Sanacora G. Riluzole in the treatment of mood and anxiety disorders. CNS drugs. 2008;22(9):761–786.

66. Zarate Jr CA, Payne JL, Quiroz J, Sporn J, Denicoff KK, Luckenbaugh D, et al. An open-label trial of riluzole in patients with treatment-resistant major depression. American Journal of Psychiatry. 2004;161(1):171–174.

67. Sanacora G, Kendell SF, Levin Y, Simen AA, Fenton LR, Coric V, et al. Preliminary evidence of riluzole efficacy in antidepressant-treated patients with residual depressive symptoms. Biological psychiatry. 2007;61(6):822–825.

68. Waubant E, Maghzi AH, Revirajan N, Spain R, Julian L, Mowry EM, et al. A randomized controlled phase II trial of riluzole in early multiple sclerosis. Annals of clinical and translational neurology. 2014;1(5):340–347.

69. Miller RG, Mitchell J, Lyon M, Moore DH. Riluzole for amyotrophic lateral sclerosis (ALS)/motor neuron disease (MND). Cochrane Database of Systematic Reviews. 2007;(1).

